# Assessing the links between pollinators and the genetic and epigenetic features of plant species with contrasting distribution ranges

**DOI:** 10.1101/2023.09.02.556048

**Authors:** Javier Valverde, Mónica Medrano, Carlos M. Herrera, Conchita Alonso

## Abstract

In flowering plants, pollinators contribute to gene flow while they also respond to variation in plant traits together determined by genetic, epigenetic and environmental sources of variation. Consequently, a correlation between abundance and diversity of pollinators and the genetic and epigenetic characteristics of plant populations such as diversity or distinctiveness is to be expected, yet no study has explored such long-term dimensions of plant-pollinator interactions. Mediterranean narrow endemics often exhibit unexpectedly high levels of population (epi)genetic diversity. We hypothesize that pollinators may contribute to explain this pattern. Specifically, we expect a stronger association of pollinators with population (epi)genetic variability in narrow endemics than in widely distributed congeners. We studied five pairs of congeneric plant species, consisting of one narrow endemic with a restricted distribution and one widespread congener, found in the Sierra de Cazorla mountains (SE Spain). In up to three populations per species, a comprehensive characterisation of pollinators was carried out to estimate pollinator diversity and visitation rate. Genetic and epigenetic diversity and distinctiveness of each population was calculated using AFLP markers and methylation-sensitive AFLP markers (MSAP) respectively. The relationships with pollinator diversity and visitation rate were assessed. The diversity of pollinators did not vary according to the plant distribution range, whereas visitation rate was higher in widespread species. As predicted, only narrow endemics showed a significant association between pollinators and their population genetic and epigenetic characteristics. Specifically, higher pollinator diversity and visitation rates entailed higher population genetic diversity and lower epigenetic distinctiveness. This work shows the value of exploring the relationships between pollinator abundance and diversity and population (epi)genetics for understanding the evolution of plant rarity.

## INTRODUCTION

Genetic diversity of animal-pollinated plant populations depends to a great extent on pollinators (Wright, 1946; Ellstrand et al. 1989; Ellstrand 1992; Ellstrand, 2003; Wessinger, 2021). In addition to the well documented effects of pollinator flight distance (e.g. Breed et al. 2015; Brunet et al. 2019; Gamba and Muchhala, 2020, 2022; Wessinger 2021), other pollinator-specific characteristics such as pollen carryover (Morris et al. 1994; Holmquist et al. 2011; Mitchell et al. 2013) or foraging strategies (i.e. number of flowers visited per plant individual; Lloyd, 1992; Eckert, 2000) determines the provenance of pollen carried by pollinators. In fact, several studies demonstrate that different pollinators with differences in the former attributes contribute with contrasting mixtures of pollen donors to the final mating portfolio of a plant individual (Castilla et al. 2017; Valverde et al. 2019). Thus, the join effect of several pollinators increases polyandry (Devaux et al. 2014; Barrett and Harder, 2017), which, in the long term could shape population genetic diversity (Pannell and Labouche 2013; Bezemer et al. 2016). On the other hand, pollinators preference towards different floral traits or plant microenvironments have been documented (e.g. Herrera, 1995; Benitez-Vieyra et al. 2006; Gómez et al. 2006, 2008; Norgate et al. 2010) suggesting that increased genetic diversity underlying large floral phenotypic variation or environmental heterogeneity could in part determine the diversity of pollinators visiting a plant population. Regardless of the direction of such relationship, there is a surprising paucity of studies exploring the correlation between the genetic diversity of plant populations and the diversity of their pollinator assemblages (see Burgin and Hopkins, 2022; Feigs et al. 2022).

In cross-talk with the genetic variation, epigenetic regulation –DNA modifications not affecting the DNA sequence– provides an additional source of potentially heritable variation. In plants, variation in epigenetic DNA methylations can predict a substantial amount of the phenotypic variance observed within natural populations (Herrera and Bazaga, 2010; Medrano et al. 2014, 2020; Schulz et al. 2014; Albaladejo et al. 2019). Significant epigenetic variation originates as a rapid response to environmental changes, and some epimutations are inherited across many generations (Richards, 2006; Turner, 2009; Verhoeven et al. 2010; Meyer, 2015; Johannes and Schmitz, 2019). These attributes confer to environmentally responsive epigenetic marks a distinct role in natural selection (Richards, 2006; Jablonka and Raz, 2009; Balao et al. 2018) and points to a potential correlation with pollinators (Alonso et al. 2019) that, as with genetic variability, may be bidirectional or simply correlative. On the one hand, due to its heritability, epigenetic variation can be expected to be subject to the same dispersal processes mediated by pollinators. On the other hand, given its ability to influence phenotype, it could influence pollinator attraction in a similar way as genetic variation. However, the versatility of epigenetic fingerprints and the difficulty of predicting the impact of gene flow on population epigenetic variability (Smithson et al. 2019; Greenspoon et al. 2022) indicate that the relationship with pollinator diversity must be complex. In combination with this, the lack of knowledge on such relationships in natural populations makes it impossible to formulate specific hypotheses, making the exploration of correlations between pollinator diversity and genetic and epigenetic variation in wild plant populations a promising avenue for understanding the ecological mechanisms underlying plant adaptation and evolution (Alonso et al. 2019).

The interrelationship between population genetic and epigenetic features and pollinator diversity may be important in explaining the adaptation of narrowly distributed plants. In theory, small and isolated populations should have reduced allelic variability due to the effects of genetic drift and inbreeding (Barret and Kohn, 1991; Ellstrand and Ellam, 1993; Gitzendanner and Soltis, 2000; Boyd et al. 2022), but genetic variability in small inbreed populations is also more sensitive to gene flow (Richards, 2000; Willi et al. 2007). In western Mediterranean mountains, many plant endemics show restricted geographical distributions consisting of patchy, small and isolated populations inhabiting harsh environments such as rocky outcrops, dolomitic soils or cliffs (Lavergne et al. 2004; Thompson et al. 2005; Molina-Venegas et al. 2015; Gimenez-Benavides et al. 2018). Several studies in these species have found higher genetic diversity than expected from the detrimental effects of drift and inbreeding (Fernández-Mazuecos et al. 2014; Jiménez-Mejías et al. 2015; Forrest et al. 2017; Medrano et al. 2020). Likewise, other studies in the same region have shown high levels of epigenetic diversity in natural plant populations (e.g. Herrera and Bazaga, 2010; Medrano et al. 2014; but see Avramidou et al. 2015) that are comparable to those found in widespread congeneric species (Medrano et al. 2020). Such findings indicate that other as yet unexplored factors may have historically influenced gene flow, counteracting the expected genetic and epigenetic impoverishment in populations of Mediterranean endemic species.

Considering that i) pollinator diversity and genetic (and potentially epigenetic) diversity may be intercorrelated, ii) the genetic (and potentially epigenetic) diversity of small, isolated populations is more sensitive to variations in gene flow, and iii) that the Mediterranean harbours a high diversity of pollinator species (Herrera, 1988; Petanidou and Ellis, 1993; Bosch et al. 1997; Herrera, 2019; Ropars et al. 2020), we hypothesize that genetic and epigenetic variation may correlate to variation in pollination diversity more obviously in Mediterranean narrow endemics than in widely distributed congeners. To test this hypothesis, we selected five pairs of congeneric species found in the Sierras de Cazorla mountain range (southern Spain), each pair consisting of one narrow endemic (restricted hereafter) and one widespread species, and analyse up to three populations per species in order to explore: 1) whether visitation rate and pollinator diversity within populations depend on the plant distribution range; and 2) if these pollinator descriptors associate with genetic and epigenetic diversities differently in narrow endemics and widespread species.

This multi-species approach and replicate population analysis at local scale takes into account phylogeny and avoids large scale environmental confounding effects (Gitzendanner and Soltis, 2000; Medrano et al. 2020) while also allows to reduce the odds of finding spurious relationships resulting from combining contemporary data (pollinator censuses) with data resulting from historical effects (genetic and epigenetic variation). Genetic and epigenetic diversities of populations are calculated from Medrano et al. (2020) and pollinator visitation rate and diversity are accurately characterised by repeated census (cf. Herrera, 2019). So far, this seems to be the first study explicitly addressing the relationships that links pollinator diversity with genetic and epigenetic features of plant populations.

## MATERIALS AND METHODS

### Site and study system

Field work was carried out in the Sierras de Cazorla, Segura y Las Villas Natural Park (Jaén Province, Spain), a mountain system which is part of the Baetic Ranges in southeastern Iberian Peninsula. This area is characterised by a complex topography and heterogeneous environmental conditions that makes it one of the richest and most diverse natural habitats in Europe (Médail and Quézel 1999; Mota et al. 2002; Melendo et al. 2003).

Five pairs of congeneric plant species (Table 1; Appendix S1) from five different families were considered, each pair consisting of one species with a restricted distribution (R, hereafter) and another species with a widespread distribution (W, hereafter). These pairs were chosen based on that they inhabit the same geographical area, and correspond to a subset of those analysed by Medrano et al. (2020): *Anthyllis ramburii* (R) and *A. vulneraria* (W); *Convolvulus boissieri* (R) and *C. arvensis* (W); *Erodium cazorlanum* (R) and *E. cicutarium* (W); *Teucrium rotundifolium* (R) and *T. similatum* (W); and *Viola cazorlensis* (R) and *V. odorata* (W). In the conceptual framework proposed by Rabinowitz (1981), all restricted species considered here have a small geographic range with small sparse populations, and with a narrow habitat specificity. Here, these species are narrow endemics from the Baetic mountain range that inhabit specific rocky microhabitats on poor sandy dolomitic soils. In contrast, widespread congeners are more common, growing on a wider spectrum of soils, usually in grasslands, shrublands or anthropised ecosystems (see Appendix S2 and Medrano et al. 2020 for a detailed description).

**Table 1.** Pollinator sampling effort. For each population of the widespread (*W*) and restricted (*R*) plant species information about the number of census performed and the average and standard deviation number of flowers per sampling patch are provided. Information on the number and probability of visits to a flower during three minutes of observation are provided. Sampling coverage is an estimate of the percentage of species sampled using the visitation to a flower as sampling unit.

| Species | Population | No. censuses | No. observed flowers per census | No. flower visits | Flower visitation prob. | Sampling coverage |
| --- | --- | --- | --- | --- | --- | --- |
| *Anthyllis ramburii* (R) | anra1 | 71 | 141.2 ± 94.5 | 90 | 0.01 | 0.97 |
|  | anra2 | 62 | 222.8 ± 207.6 | 901 | 0.05 | 1.00 |
|  | anra3 | 65 | 60.2 ± 39.7 | 36 | 0.02 | 1.00 |
| *Anthyllis vulneraria* (W) | anvu1 | 62 | 200.3 ± 93.7 | 500 | 0.05 | 0.99 |
|  | anvu3 | 92 | 317.7 ± 199.5 | 505 | 0.02 | 0.99 |
| *Convolvulus boissieri* (R) | cboi2 | 77 | 19.4 ± 21.5 | 125 | 0.1 | 0.99 |
|  | cboi3 | 72 | 23.1 ± 9.9 | 137 | 0.12 | 0.96 |
| *Convolvulus arvensis* (W) | carv1 | 69 | 5.1 ± 1.5 | 82 | 0.24 | 0.92 |
|  | carv2 | 68 | 9.0 ± 3.1 | 149 | 0.4 | 0.94 |
| *Erodium cazorlanum* (R) | ecazF | 77 | 5.4 ± 3.8 | 29 | 0.08 | 0.76 |
|  | ecazL | 72 | 11.9 ± 5.3 | 60 | 0.08 | 0.90 |
|  | ecazT | 37 | 2.6 ± 1.0 | 21 | 0.19 | 0.91 |
| *Erodium cicutarium* (W) | ecicC | 101 | 21.1 ± 13.6 | 115 | 0.06 | 0.92 |
|  | ecicF | 86 | 8.8 ± 4.8 | 59 | 0.06 | 0.95 |
|  | ecicT | 87 | 3.3 ± 1.4 | 42 | 0.15 | 0.79 |
| *Tecucrium rotundifolium* (R) | trot1 | 71 | 57.8 ± 35.5 | 104 | 0.03 | 1.00 |
|  | trot2 | 66 | 38.8 ± 20.4 | 176 | 0.07 | 0.99 |
|  | trot3 | 62 | 58.1 ± 42.7 | 1706 | 0.54 | 1.00 |
| *Tecucrium similitum* (W) | tpol1 | 66 | 252.6 ± 118.1 | 1028 | 0.06 | 1.00 |
|  | tpol2 | 60 | 122.0 ± 79.0 | 574 | 0.06 | 1.00 |
|  | tpol3 | 60 | 232.6 ± 111.0 | 2947 | 0.21 | 1.00 |
| *Viola cazorlensis* (R) | vcaz1 | 63 | 21.7 ± 13.4 | 38 | 0.05 | 0.98 |
|  | vcaz2 | 65 | 49.1 ± 34.4 | 81 | 0.02 | 1.00 |
|  | vcaz3 | 70 | 22.9 ± 15.1 | 20 | 0.02 | 1.00 |
| *Viola odorata* (W) | vodo1 | 107 | 22.0 ± 12.7 | 25 | 0.01 | 1.00 |
|  | vodo2 | 74 | 32.3 ± 23.7 | 35 | 0.01 | 1.00 |

### Sampling and data acquisition

#### Pollinator data

We initially selected three populations per species (Medrano et al. 2020). However, herbivory damage, insufficient flowering individuals or logistic limitations limited our study to only 26 populations. During March-June 2021 the pollinator assemblage of each population was characterised by conducting a minimum of 60 pollinator censuses. Here, a pollinator census is the sampling unit and consisted of 3-min of observation of a flowering patch, i.e. a delimited area with a number of flowers that are easily accessible to visual inspection. At each census, the identity of each pollinator and the number of flowers visited, as well as the number of open flowers in the patch, were recorded. Censuses were carried out in randomly selected patches, always in direct sunlight and on sunny days and were distributed at the beginning and at the peak of flowering to capture a fair representation of the pollinator fauna (Valverde et al. 2014). This sampling method and observation effort have demonstrated to give reliable estimates of pollinator diversity in a wide range of plant species (Herrera, 2019, 2020).

In this study, a pollinator is considered to be any taxon that at any time contacted the reproductive organs of the plant. For a precise taxonomical identification of pollinators, each pollinator-flower interaction was photographed. Difficult to identify specimens were collected and contrasted against our own reference pollinator checklist (Ortiz et al. 2023 unpub.) or sent to a number of experts listed in the acknowledgements for identification. Because of the unavoidable uncertainty in the identification of certain taxonomical groups, some specimens were clumped. For example, *Lasioglossum pauperatum* also includes *L. transitorium*, Nitidulidae species were all considered *Brassicogethes aeneus* as this species represents 75% of all specimens in most of these plant species (Herrera and Otero, 2021). In other cases, we used morphospecies as for example for specimens from the Tachinidae or Anthicidae families or from the *Empis* or *Hilara* genus (Empididae) (Appendix S4 resumes all observed species). As a result, up to 92% of pollinator individuals were characterised at the species or morphospecies level.

It is important to highlight that, the availability of pollinator data from previous years of some of our study populations (Herrera 2018, 2019) allowed us to compare our census data with data collected 5 to 16 years before with this same methodology (see Appendix S3 for a detailed description and results of these analyses). This comparison showed a high correlation in species richness between years (*β* = 0.80, *t* = 4.15, df = 10, *P* = 0.002). This finding supports the interannual stability of pollinator diversity in our study system as previously shown in Herrera (2018) for a much larger number of species in the study area, and allows us to be more confident when comparing contemporary pollinator data with historically inherited (epi)genetic data of plants.

#### Genetic and epigenetic data

The genetic and epigenetic data are a subset of those published in Medrano et al. (2020). Specifically, from the populations for which pollination data were obtained, an average of 26 random individuals per population were genotyped (range 23–40). Genetic profiles of sampled individuals were obtained by amplified fragment length polymorphism (AFLP; Meudt and Clarke 2007) and epigenetic profiles by amplified methylation-sensitive polymorphism (MSAP; Reyna-López et al. 1997; Fulneček and Kovařík, 2014; Schulz et al. 2014), a technique useful to identify genome-wide methylation profiles in ecological epigenetics studies of species without detailed genomic information (Schrey et al. 2013). As a result, for the AFLP markers we obtained a binary matrix depicting the presence or absence of each loci. For the MSAP markers, we followed Schulz et al. (2014) to obtain two binary matrices: one with hemi– or fully methylated epiloci (hereafter M-MSAP), and one data matrix of unmethylated epiloci (U-MSAP). We eliminated inconsistent loci after measuring their repeatability in a subset of samples (8.6–17.6 % of samples for AFLP; 13.3– 29.3% of samples for MSAP). We also discarded non-informative monomorphic loci by selecting those in which the proportion of presence or absence of the fragment exceeded 5%. This yielded 140 to 316 AFLP, 84 to 181 U-MSAP and 100 to 213 M-MSAP loci per species. For a detailed description of these methodologies please refer to Appendix S5 or to Medrano et al. (2020).

### Data analyses

#### Pollinator descriptors

The pollination regime of each population was characterised by the visitation rate and pollinator diversity. We calculated the visitation rate using two indices: the patch visitation probability (visit × 3-min census^-1^), which is irrespective of the number of flowers visited, and the flower visitation probability (visited flowers × no. open flowers^-1^ × 3-min census^-1^), which takes into account the number of flowers visited.

We estimated the sample coverage of each population (proportion of the total estimated number of species; Chao and Jost, 2012) based on the number of visited flowers and using the *R* package *iNEXT* (Hsieh et al. 2016). Following, we calculated the diversity of pollinators on each population using Hill numbers (*^q^D*; Hill, 1973) at an equal sample coverage of 0.8. At this value, the extrapolation of diversity values of the two populations showing lower sampling coverages (ecazF, 0.76 and ecicT, 0.79; Table 1) is reliable (Chao et al. 2014).

Hill numbers are a unified family of diversity indexes that equals the number of equally abundant taxa needed to obtain a given diversity, allowing for easier comparisons among assemblages (Jost, 2007). These are defined as:

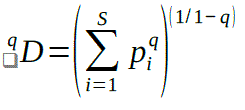

Where *S* is the total number of species, *p_i_* is the relative frequency of species *i*, and *q* is a parameter that determines the sensitivity of the index to the relative frequencies. By varying *q* between 0 and 2, Hill numbers range between species richness (*q* = 0), the logarithm of the Shannon information index (*q* ≈ 1, Shannon’s diversity herein) which considers true relative abundances of species, and Simpson’s diversity (*q* = 2) which weights for dominant species (Hill, 1973; Jost, 2006; Chao et al. 2014). To obtain a fine characterisation of the pollinator diversity of each population, we constructed the diversity profiles by calculating the Hill numbers from *q* = 0 to *q* = 2 by increments of 0.1, allowing us to visually explore the diversity along a gradient of relative importance of pollinator abundances (Jost, 2007).

#### Genetic and epigenetic descriptors

Each population was characterised by its genetic and epigenetic diversity and distinctiveness. We calculated the diversity for the genetic and epigenetic markers by means of two indices: the Proportion of Polymorphic Loci and the per-locus average of the Shannon diversity index (King and Schaal, 1989; *R* package *MSAP*, Pérez-Figueroa, 2013; Appendix S6). To calculate the genetic and epigenetic distinctiveness, we used two indexes: the Proportion of Private Loci and a Rarity Index (Schönswetter and Tribsch, 2005; *R* package *AFLPdat*, Ehrich, 2006; Appendix S6). Values from the Rarity Index are expected to be high at isolated populations with an historical accumulation of rare markers.

#### Statistical analysis

Differences in pollinator diversity between restricted and widespread species were analysed using species richness, Shannon’s and Simpson’s indexes (Hill numbers of q equal to 0, 1 and 2, respectively). For each diversity index, we calculated the difference between each pair of populations from the restricted and the widespread congeners and used non-overlapping confidence intervals as criteria to assess the significance of each comparison.

Differences in visitation rate between restricted and widespread species were assessed using generalized mixed-effects models (*R* package *lme4*, Bates et al. 2015). We modeled the global effect of species distribution type across all congeneric pairs of species using a binomial error distribution and a log-link function. In these models, population was nested within genus as random effects. Further, we assessed the effect of species distribution at each congeneric pair through additional models that included population as random effects and genus as a fixed term. Such models were applied using both the patch and the flower visitation probabilities as response variables. For the patch visitation probability the scaled number of flowers per species was included as a covariate to control for the potential effect of flower display in attracting pollinators. For the flower visitation probability, an observation-level random effect was included to deal with overdispersion in our data (Harrison, 2014). Differences between widespread and restricted species at each model were assessed through *post-hoc* pairwise contrasts on the estimated marginal means and using Bonferroni correction in models considering multiple comparisons (*R* package *emmeans* V.1.5.4, Lenth, 2021).

Finally, we assessed the relationship between pollinators and the (epi)genetic diversity and distinctiveness of populations. First, we used Pearson correlations between (epi)genetic and pollinator parameters to explore general trends in populations from restricted and widespread species. Correlations with Hill diversities of intermediate q values were included in this exploration to visualize such relationships along a gradient of relative importance of pollinator abundances.

Following, a more exhaustive analysis was performed using generalized mixed-effects models on the relationships showing a significant correlation. By including species as a random term, these models allowed to control for variation among species. Family and linkage functions were chosen in each model according to data structure and goodness-of-fit (see details at Results section). For all relationships, the genetic and epigenetic parameters were used as dependent variables and plant genus was included as random factor. In all cases we used random intercept models because of our limited data per species and because these outperformed in all cases the random slopes models (Appendix S7).

## RESULTS

### Pollinator composition

We recorded 1214 pollinator individuals from 179 different taxa. The relative frequency of pollinator orders varied considerably across sample populations and plant species (Fig. 1). Hymenoptera performed 84.1% of all visits, but were not the most frequent pollinators in all study species (Fig. 1 and Appendix S8). At two genus [*Anthyllis* and *Teucrium*] congeneric species showed similarities in the main pollinator order. In *Anthyllis* species Hymenoptera were the main pollinators (*A. vulneraria* (*W*) = 85–94% of all flower visits; *A. ramburii* (*R*) = 71–100%), although they differed in the identity of the main pollinator family: Apidae (79–85%) –mainly Anthophorini– in *A. vulneraria* and Andrenidae (44–72%) and Apidae (53%) in *A. ramburii*. In *Teucrium*, Hymenoptera were the main pollinators in all populations from the widespread *T. similatum* (*W*; 78–98%) and in two populations of the restricted *T. rotundifolium* (*R*; 73–94%), while Diptera were the most frequent pollinators in the third population (43%). On the other hand, contrasting pollinator assemblages were found between congeneric species in *Convolvulus*, *Erodium* and *Viola* genera. In *Convolvulus*, Coleoptera were the main pollinators in *C. arvensis* (*W*; 51–59%) while Hymenoptera dominated in *C. boissieri* (*R*; 52–67%). In *Erodium*, Diptera visited most flowers in *E. cicutarium* (*W*; 50–83%), while in *E. cazorlanum* (*R*) Hymenoptera were the main flower visitors in two populations (44–53%) and Coleoptera in the other (71%). Finally, in *Viola odorata* (*W*), Hymenoptera –mainly *Anthophora dispar*–were the main pollinators (76–100%), while in *V. cazorlensis* (*R*) Diptera –mainly Bombyliidae– were the main pollinators in two populations (55– 84%) and Lepidoptera –mostly *Macroglossum stellatarum* (Sphingidae)– in the other (62%).

**Figure 1.**
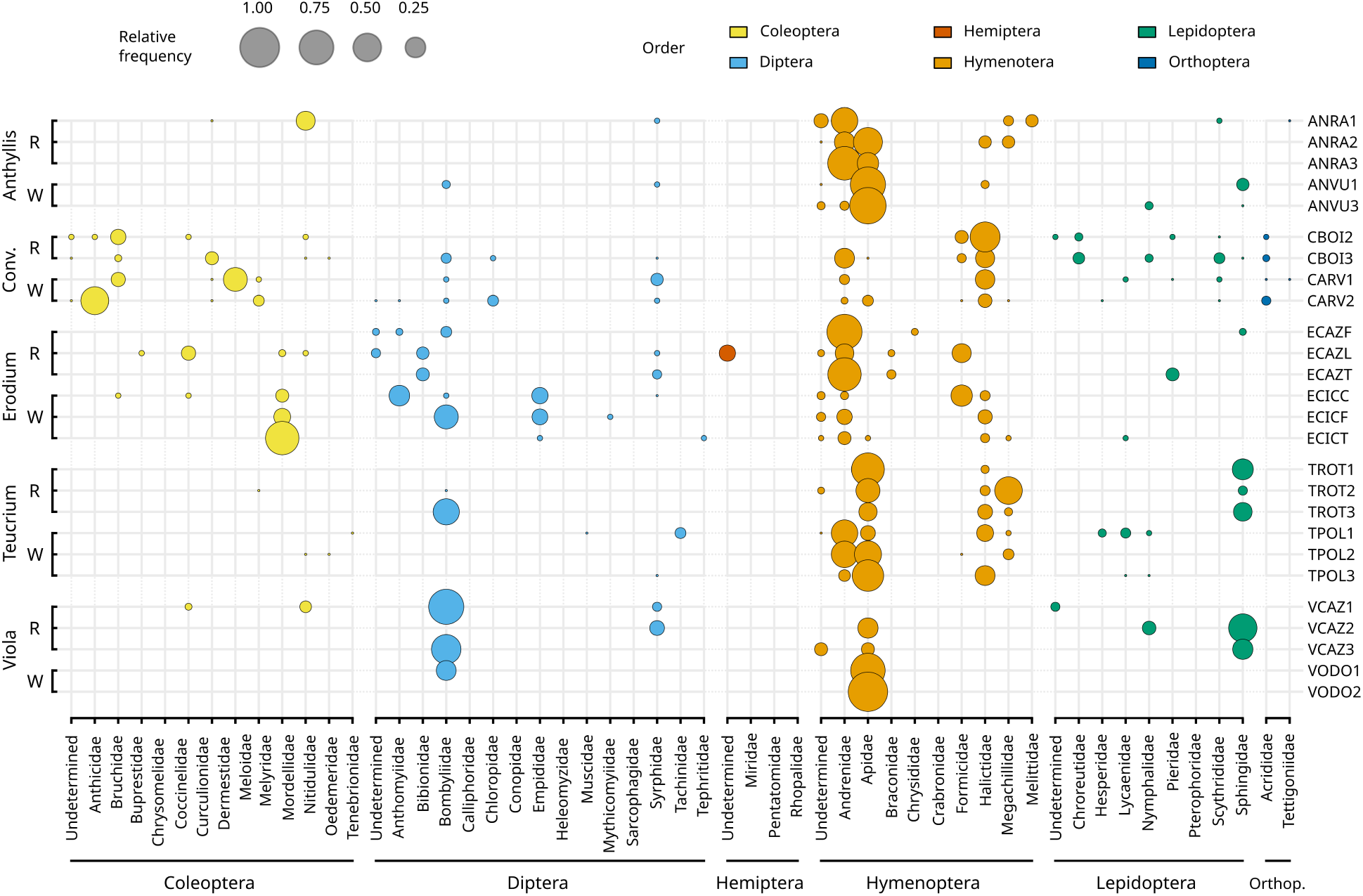
Pollinator assemblages observed at study populations. Circle sizes are relative to the proportion of flower visits made by each insect family in a given population. *R* and *W* stands for restricted and widespread species respectively. *Conv.* denotes *Convolvulus*, and *Orthop.* Orthoptera.

### Variation in pollinator diversity and visitation rate

The diversity profiles did not suggest any consistent trend in the comparison of pollinator diversities between populations of restricted and widespread congeneric species (Appendix S9). Only two genera showed consistent differences in pollinator diversity between the restricted and the widespread species (Fig. 2). In *Anthyllis*, flowers of the widespread *A. vulneraria* were visited by a more diverse pollinator assemblage than flowers from its restricted congener, with non-overlapping confidence intervals in most comparisons of the pollinator richness, Shannon and Simpson diversity (Fig. 2). In contrast, the restricted *Viola cazorlensis* showed significantly higher pollinator diversity in most comparisons. Apart from these two exceptions, the rest of congeneric pairs did not show any clear difference in pollinator diversity of widespread and restricted species.

**Figure 2.**
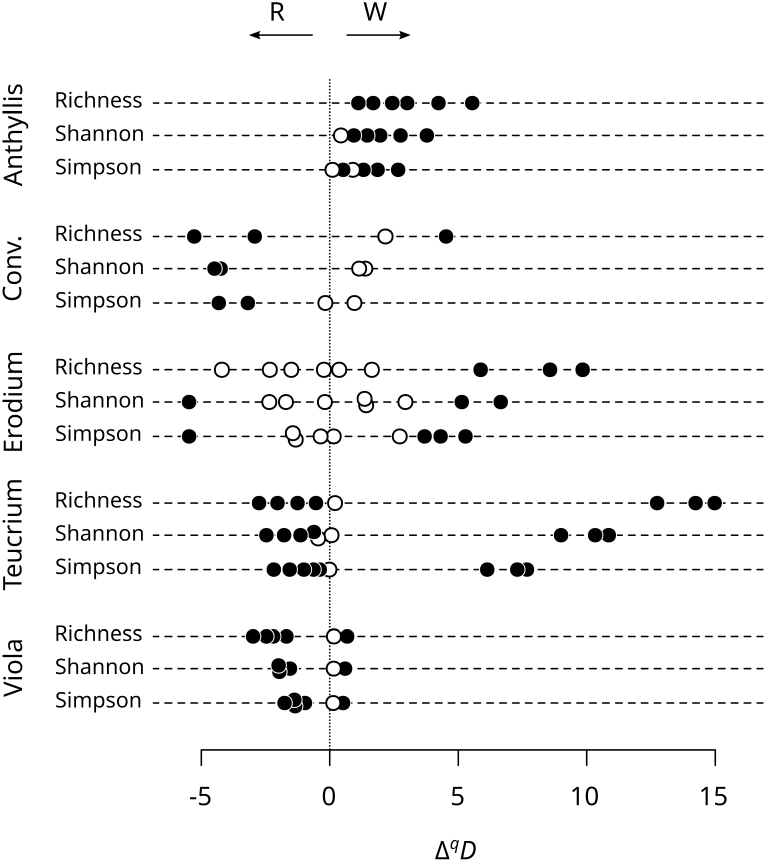
Differences in pollinator diversity between congeneric populations. Each point represents the difference in diversity between a population from a widespread plant species and a population from the corresponding restricted congener (*Δ^q^D*). Thus, negative values indicate higher diversity in the population from restricted species (*R*) while positive values indicate higher diversity at the population from the widespread congener (*W*). Filled symbols denote comparisons with non-overlaping confidence intervals

On the contrary, the type of distribution range had an effect on the visitation rate. The patch visitation probability varied with the distribution range (estimate = –0.73, *z* = –2.08, *P* = 0.037; Appendix S10), being higher for widespread species (Fig. 3). Flower display showed a positive significant effect on the patch visitation probability (estimate = 0.33, *z* = 5.37, *P* < 0.001). The higher patch visitation probability in the widespread species was particularly noticeable in the *Teucrium* pair (*z*_ratio_ = 4.28, *P* < 0.001; Fig. 3; Appendix S11). The flower visitation probability was also influenced by the distribution range (estimate = –0.74, *z* = –1.95, *P* = 0.05; Appendix S10), with flowers from the widespread species having higher visitation probabilities (Fig. 3). This global pattern reflected the significantly higher probabilities found for the widespread species in the genera *Anthyllis*, *Convolvulus* and *Teucrium* (*z*_ratio_ = 1.97–6.08, *P* < 0.05; Fig. 3; Appendix S11).

**Figure 3.**
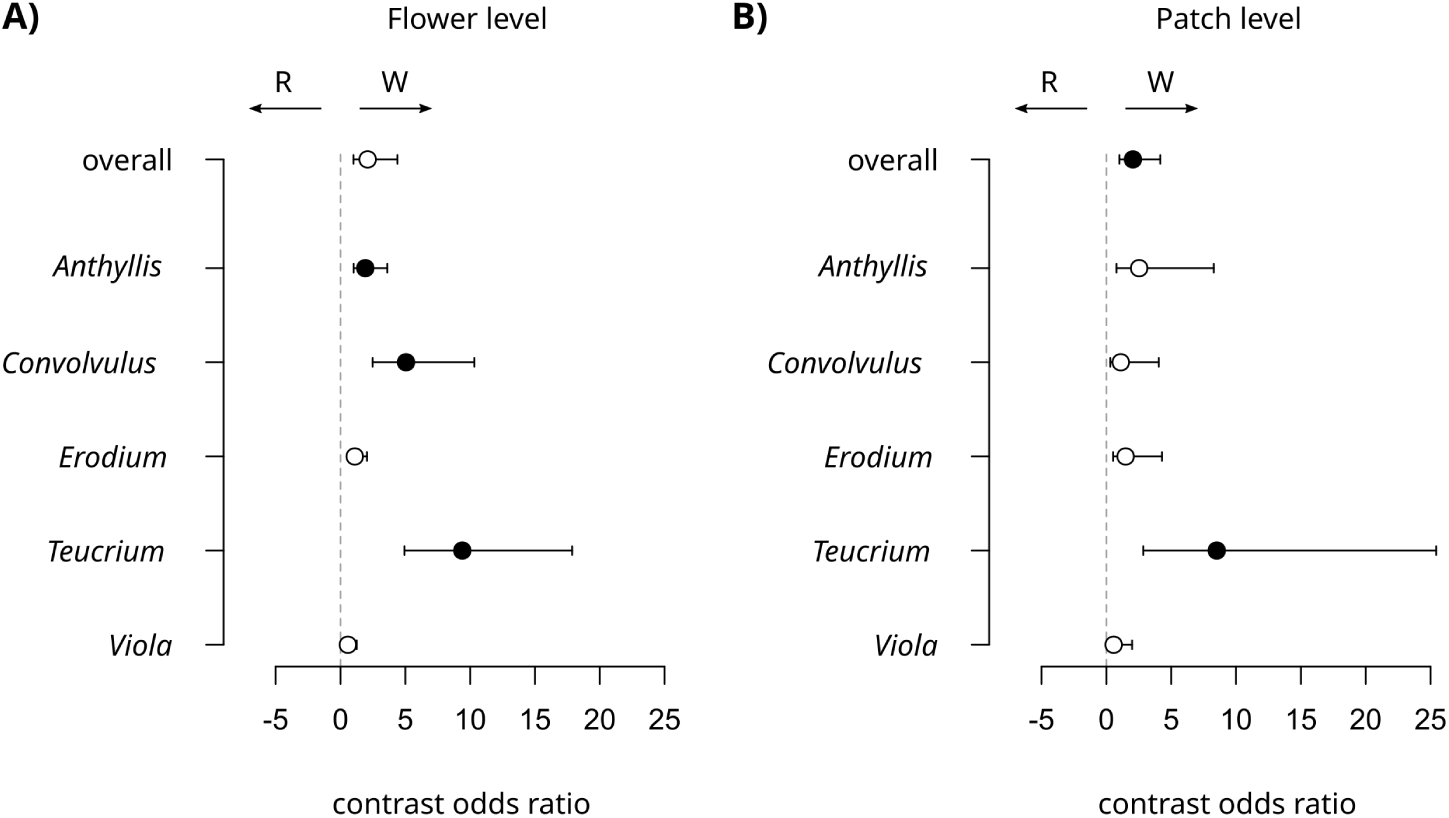
Differences in visitation probabilities between congeneric populations. Panels show posthoc paired comparisons of visitation probability at the patch (A) and flower (B) levels. Estimated odds ratios and the corresponding 95% confidence interval are shown for each comparison. Negative and positive values indicate higher visitation rates in the restricted and widespread species respectively. Filled symbols denote significant differences.

### Relationship between pollinators and plant population genetics and epigenetics

Only the restricted species showed significant correlations between pollinators and their genetic or epigenetic population descriptors (Appendix S12). Regarding the (epi)genetic diversity, the Shannon diversity of AFLP genetic markers showed positive significant correlations with patch and flower visitation rates (*β* = 0.59, *t* = 2.53, df = 12, *P* = 0.026 and *β* = 0.54, *t* = 2.19, df = 12, *P* = 0.049, respectively) and with pollinator richness (*β* = 0.60, *t* = 2.58, df = 12, *P* = 0.24). Similarly, the Shannon diversity of U-MSAP epigenetic markers showed a positive significant correlation with the patch visitation probability (*β* = 0.59, *t* = 2.51, df = 12, *P* = 0.027) and with pollinator richness (*β* = 0.57, *t* = 2.40, df = 12, *P* = 0.034) and pollinator Shannon’s diversity (*β* = 0.53, *t* = 2.19, df = 12, *P* = 0.044). After controlling for interspecies variability, the mixed-effects models showed a significant relationship between AFLP Shannon diversity and patch visit probability (estimate = 0.08, *t* = 2.59, *p* = 0.027; Table 2), while a decrease in significance was observed for the rest of the previous significant correlations (*P* > 0.105).

**Table 2.** Summary of the significant relationships observed between pollinator and (epi)genetic descriptors in plant species with a restricted distribution range. The table shows the correlations and mixed model results of those relationships that are significantly correlated (see figures 4 and S8). Note that for the proportion of privative methylated epialleles (M-MSAP) the coefficients from the mixed effects models correspond to estimates using the log-link function. *P*-values in bold denote significance at α = 0.05.

| Descriptors of (epi)genetic diversity |  | Pearson's correlation |  |  | Linear mixed effects model |  |  |
| --- | --- | --- | --- | --- | --- | --- | --- |
| Response | Predictor | Correlation | *t* | *p* | Estimate | *t* | *P* |
| AFLP Shannon index | *Visitation probability (patch)* | 0.59 | 2.53 | **0.026** | 0.08 | 2.59 | **0.027** |
|  | *Visitation probability (flower)* | 0.54 | 2.19 | **0.049** | -0.04 | -0.21 | 0.837 |
|  | *Pollinator richness (<sup>0</sup>D)* | 0.60 | 2.58 | **0.024** | 0.00 | 1.27 | 0.234 |
| U-MSAP Shannon index | *Visitation probability (patch)* | 0.59 | 2.51 | **0.027** | 0.08 | 1.76 | 0.109 |
|  | *Pollinator richness (<sup>0</sup>D)* | 0.57 | 2.39 | **0.034** | 0.00 | 1.78 | 0.105 |
|  | *Pollinator Shannon diversity* | 0.53 | 2.19 | **0.048** | 0.00 | 1.56 | 0.149 |
| Descriptors of (epi)genetic distinctiveness |  | Pearson's correlation |  |  | Linear mixed effects model |  |  |
| Response | Predictor | Correlation | *t* | *p* | Estimate | *t* | *P* |
| AFLP Rarity index | *Prob. patch visit* | -0.54 | -2.20 | **0.048** | -1.01 | -1.44 | 0.180 |
| M-MSAP Proportion of privative alleles (log-link) | *Pollinator richness* | -0.55 | -2.31 | **0.039** | -0.47 | -6.42 | **<0.001** |
|  | *Pollinator Shannon diversity* | -0.57 | -2.39 | **0.034** | -0.59 | -6.89 | **<0.001** |
|  | *Pollinator Simpson diversity* | -0.55 | -2.28 | **0.041** | -0.76 | -6.93 | **<0.001** |

Models use the following syntaxes in the *lme4* R package: *lmer*( $Y \sim X + (1 | \text{genus})$ ) and *glmer*( $Y \sim X + (1 | \text{genus})$ , *family = gaussian(link="log")*) for the M-MSAP proportion of privative alleles. In these models Y and X denote the response and predictor variables respectively.

As for the (epi)genetic distinctiveness, the rarity index of AFLP genetic markers showed a negative significant correlation with the patch visitation probability (*β* = –0.54, *t* = –2.20, df = 12, *P* = 0.048), while the proportion of M-MSAP epigenetic privative markers had a negative significant correlation with all the range of pollinator diversities (*β* = –0.57– –0.55, *t* = –2.28– –2.41, df = 12, *P* < 0.042; Fig. 4 and Appendix S12). Mixed-effect models showed a significant negative relationship of the proportion of M-MSAP with any measurement of pollinator diversity using a log-link function (*β* = –0.47– –0.76, *t* = –6.42– –6.94*, P* < 0.001; Table 2; Fig. 5).

**Figure 4.**
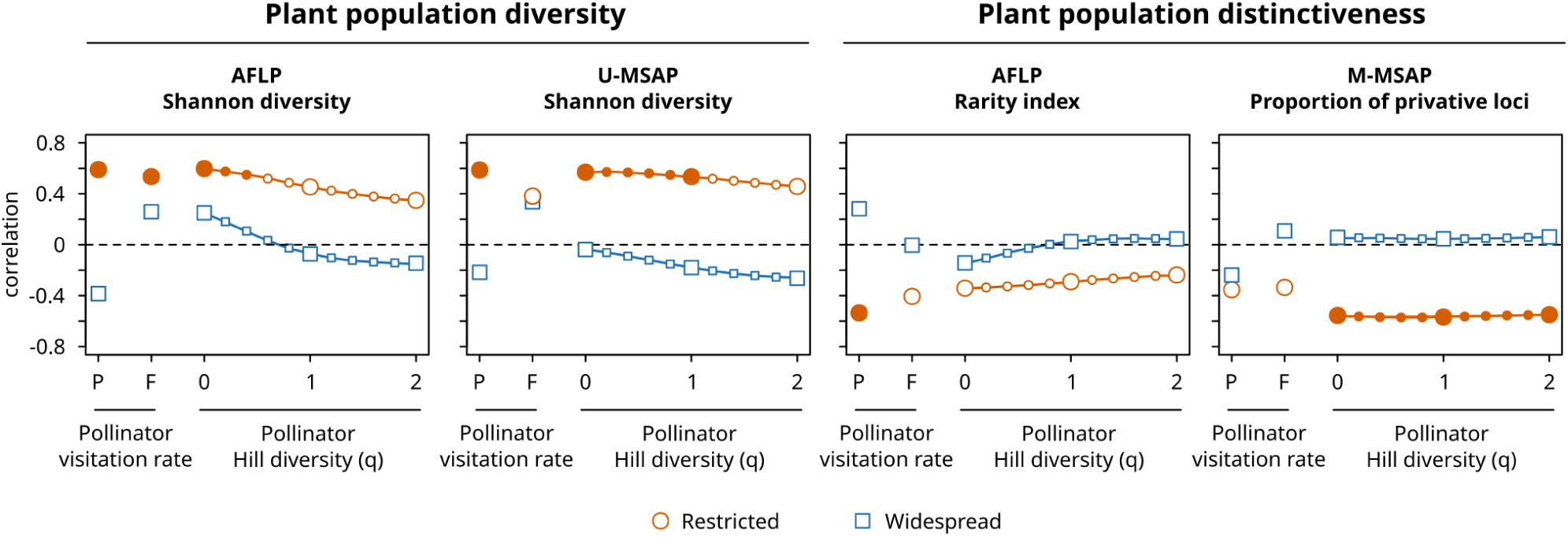
Correlations between estimators of pollinator and genetic and epigenetic descriptors. The panels show the correlations for the diversity and distinctiveness indices for which any significant correlations were found (for all correlations see figure S8). Orange circles depict restricted species and blue squares widespread species. Filled symbols indicate significant correlations at α = 0.05. Increased size symbols highlight correlations at the species richness, Shannon and Simpson diversities (*q* equal to 0, 1 and 2 respectively). See Material and Methods for an accurate description of these indices.

**Figure 5.**
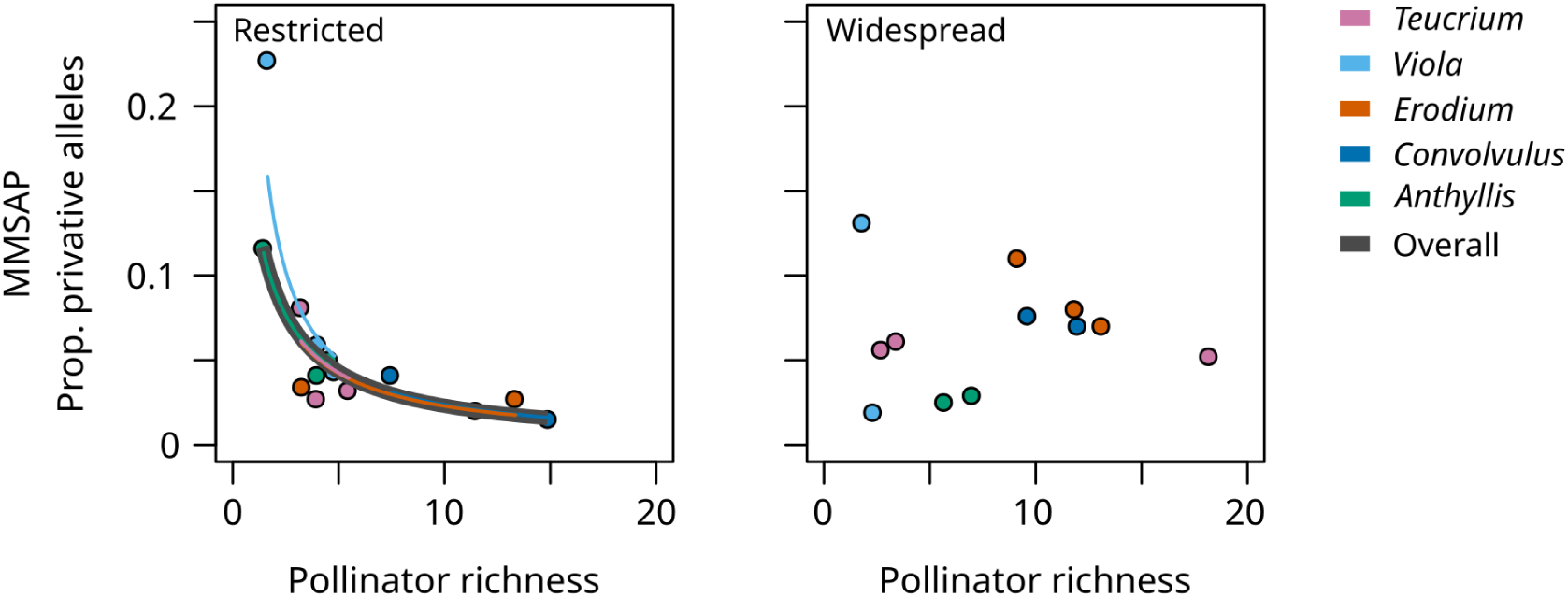
Relationship between the proportion of methylated privative alleles and pollinator richness. Model fit for the restricted species is shown in the left panel: predicted relationships for all populations (fixed effect level; thick grey line) and for each plant species separately (random effect levels; thin coloured lines) are depicted.

## DISCUSSION

This study explores whether genetic and epigenetic variation in plant populations correlate to variation in pollinator diversity and abundance more obviously in narrow endemics than in widely distributed congeners. The study undertakes what seems to be the first research that explicitly addresses the relationships linking pollinator diversity and visitation rate with genetic and epigenetic features of plant populations.

The data obtained operate at two temporal levels. On the one hand, pollinator data represent a contemporary picture of plant-pollinator interactions. On the other, genetic and epigenetic data reflect the historical accumulation of evolutionary processes in the same study populations. Ecological data must therefore be sufficiently representative of such a temporal process if reliable conclusions are to be drawn given that in many cases plant populations may undergo interannual turnover in their pollinators (e.g. *Lavandula latifolia*, Herrera,1988; *Erysimum mediohispanicum*, Valverde et al. 2014). In this study, plants show temporal stability between years in both visitation rate (see Herrera, 2019) and pollinator richness (see analysis in Appendix S3), indicating that the current pollinator data are sufficiently representative of past situations, which provides some confidence for comparison with population genetic data. Thus, although interpreted with caution, the results of this study provides insights on the pollinator-(epi)genetic relationships that allows a better understanding of the ecology of narrow endemic species.

In the following sections we first provide a general, naturalistic overview of the observed pollinator diversity. We continue by discussing the differences in pollinator diversity and visitation rates between restricted and widespread species. Finally, we discuss the relationship between the diversity of pollinators and the diversity of plant genetic and epigenetic markers.

### Pollinator composition

We recorded at least 179 pollinator species in the 26 populations studied. This value is in consonance with previous studies reporting the high diversity of pollinator species found in these mountains (Herrera, 2019, 2021) and adds to previous evidences that the Mediterranean region is a hotspot of pollinator diversity (e.g. C. M. Herrera, 1988, 2019; J. Herrera, 1988; Petanidou and Ellis, 1993; Bosch et al. 1997; Ropars et al. 2020). Hymenoptera species were the most important visitors in most plant species and populations (84% of flower visits). Although the importance of this group for plant reproduction is well-acknowledged (Ollerton, 2017; Patel et al. 2021), other historically underestimated taxonomic groups may be also important pollinators (e.g. Jaukers and Wolters, 2008; Orford et al. 2015; Valverde et al. 2019). Here, the variability in ranked frequency of pollinator visits evidences the relevance of non-hymenoptera species (see Appendix S8). For instance, Coleoptera, Diptera or Lepidoptera, stood out as the most frequent visitors to flowers of some plant species (e.g. *C. arvensis*, *V. cazorlensis*) or in certain populations, evidencing a high number of non-hymenopteran pollinators and highlighting the need to include all taxa in pollinator surveys in order to assess the true diversity of pollinators.

Pollinators have been posed as key in the evolution of rarity in flowering plants (Orians, 1997), yet the few studies that analyse this prediction show discrepancies in their results, making it difficult to draw general conclusions. This is probably due to ambiguity in classifying species as rare (Rabinowitz, 1981) and the way studies are conducted –e.g. single species studies (Jabis et al. 2011; Fernández et al. 2015) versus comparative approaches (Merhoff, 1983; Karron, 1987; Purdy et al. 1994) –, but also to the diverse features of the study species (Lavergne et al. 2004; Giatzouzaki et al. 2022). Aiming to overcome these problems, our study considers several pairs of restricted and widely distributed congeneric species (according to Rabinowitz, 1981) from different plant families. As a consequence, study species display contrasting floral characteristics that may partly explain the observed differences in pollinator composition. For instance, plant species bearing flowers with a restricted access to nectar were mainly visited by few insect orders or families (e.g. Lepidoptera and Bombyliidae in *V. cazorlensis*, Hymenoptera in *V. odorata*, and in *Teucrium* and *Anthyllis*). While species with more accessible flower morphologies (i.e. *Erodium*, *Convolvulus*) were visited by a larger number of pollinator orders or families. Furthermore, the flowering periods of the species studied are spread over four months encompassing spring, so it is expected certain seasonality in pollinator composition resulting from insect phenologies (Herrera et al. 2022). For example, while in the earliest flowering species *V. odorata* the diversity of pollinators is the lowest recorded, the latest flowering *Convolvulus* species are among the most diverse at different taxonomical levels. As in previous studies, heterogeneity in the floral features and flowering time of the study species is likely to influence contrasting responses between the congeneric pairs in pollinator diversity and visitation rate. However, by considering them all in a multispecies population framework allows to explore more generalisable patterns discussed below.

### Variation in pollinator diversity and visitation rate

Our results support previous findings on the differences between restricted and widespread congeners in the diversity and visitation rates of pollinators. First, we did not find a clear effect of distribution type on pollinator diversity. Consistent with the study by Karron (1987), three of the congeneric pairs studied showed no clear pattern in diversity differences, while the other two pairs showed clear but contrasting differences. In *Anthyllis*, the widespread *A. vulneraria* showed higher species diversity than its restricted congener in most comparisons, although it has to be noted that at higher insect taxonomical levels (e.g. family, order) the restricted *A. ramburii* was visited by a higher number of taxa indicating a more diverse functional diversity. On the contrary, in the *Viola* genus the restricted *V. cazorlensis* showed higher diversity in most comparisons. Very likely, the latter result reflects the contrasting flowering periods of these two species. Specifically, the earlier flowering of the widespread congener could be a limiting factor for the availability of pollinator species.

Second, we observed that both patches and individual flowers of widespread species showed a tendency towards higher visitation rates, a result that is consistent with previous findings on this issue (Merhoff, 1983; Karron, 1987; Purdy et al. 1994; Rymer et al. 2005; Powell et al. 2011; but see Banks, 1980 and Ruilova and Marques, 2016). This pattern was particularly clear in three congeneric pairs: *Anthyllis*, *Convolvulus* and *Teucrium* pairs (Appendix S11). The widespread species from two of these pairs with restricted corollas (*A. vulneraria* and *T. similatum*) displayed considerably more flowers per patch than their respective restricted congeners (Table 1) suggesting a positive effect of flower display in attracting more pollinator visits (e.g. Willson and Price, 1977; Thomson, 1988; Klinkhamer et al. 1989; reviewed in Ohashi and Yahara, 2001). In fact, the observed positive and significant effect of the number of flowers on the patch visitation probability would corroborate this. Other uncontrolled variables such local soil substrate and its effect on pollinator abundance (Potts and Willmer, 1997; Potts et al. 2005; Sardiñas and Kremen, 2014) may be responsible of the observed differences, specially given that all study species inhabit rocky outcrops with poor soils. Future studies addressing differences between restricted and widespread species should include restricted species inhabiting other soil types to corroborate this latter explanation.

### Relationships between pollinators and plant population genetics and epigenetics

According to our predictions, in this study plant population genetic and epigenetic variation correlated to variation in pollination diversity more obviously in restricted species than in widespread congeners. As for the (epi)genetic diversity, our analyses showed that higher visitation rates and pollinator diversity related to higher genetic and epigenetic diversity of populations from restricted plant species. Such relationship was similar for the three markers, but significant for the genetic (AFLP) and the unmethylated epigenetic markers (U-MSAP). This finding may be related to the smaller population sizes that by definition differentiate restricted from widespread plant species (Rabinowitz, 1981). For instance, in small populations, correlated paternity –the proportion of full siblings within maternal progeny arrays– is higher (Hardy et al. 2004), thus promoting inbreeding and the consequent reduction in genetic diversity (Wright, 1969; Ellstrand and Elam, 1993). Under these circumstances, higher pollinator diversities could counteract these expected outcomes by the deposition of more genetically diverse pollen loads on stigmas (Castilla et al. 2017; Rhodes et al. 2017; Valverde et al. 2019). Alternatively, given that population genetic and epigenetic diversity can reflect phenotypic variability (Gómez et al. 2009; Herrera and Bazaga 2009; Wilschut et al. 2016) or environmental heterogeneity (Richardson et al. 2014; Valverde et al. 2016), the positive relationship found between pollinator and (epi)genetic diversity could reflect the effect of higher phenotypic variability or higher environmental heterogeneity in attracting more pollinators (Herrera, 1995; Gómez et al. 2008). Clarifying any or both of these hypotheses requires studies with more and better characterised populations. In particular, measures of the genetic or sire diversity of the progeny of individual plants and a fine characterisation of the phenotypic and microenvironmental variability within populations can be key to assert the causal direction of the observed relationship. On top of this, the observed decrease in significance when applying mixed-effects models indicates that the magnitude of the effect is highly dependent on the species studied. Thus, further advances in this issue should focus in single species or narrow the similarities in life traits between the studied species to minimize further sources of variability.

As for the (epi)genetic distinctiveness, we found a negative association between methylated epigenetic distinctiveness and pollinator diversity again, only at the restricted species. In contrast to the previously discussed relationship, this correlation was consistent along the whole range of the explored pollinator Hill diversities and maintained even after taking into account the variation between plant species. Regardless of the effect on plant fitness, eventual immigrant genetic (and perhaps epigenetic) alleles may spread more rapidly in small populations than in large populations, especially if the inbreeding level of a population is high (reviewed in Tallmon et al, 2004 and Whiteley et al. 2015). This suggests that populations of rare species may be more sensitive to seed or pollen flow than populations from widespread species. In addition, several studies have experimentally demonstrated that the proportion of immigrant pollen flow may be higher in small populations (eg. Klinger et al. 1992; Goodell et al. 1997). In this sense, the observed negative linear relationship between epigenetic divergence and pollinator diversity in restricted species could be related to a greater effect of pollen flow between populations associated with higher pollinator diversity. For instance, the relationship between pollinators and the genetic cohesion of populations has been studied mainly by comparing plant species or populations whose main pollinators widely differ in their flight capabilities (e.g. Castellanos et al. 2003; Breed et al. 2015; Gamba and Muchhala 2020 and references therein). However, other studies have also found evidences of pollen flow between populations pollinated my smaller insects with presumably lower flight distances (e.g. Dick et al. 2003; Byrne et al. 2008). Thus, despite the probability of pollen transport between populations by a single pollinator is very low, the integration of a higher diversity of pollinator flight kernels may make such events eventually relevant. This potential interpretation must be considered with caution and understood under the assumption that in our study species, pollinator diversity is relatively stable over the years. However, our results should serve as a correlative evidence for future studies to consider pollinator diversity as a potential factor affecting genetic cohesion between populations.

The fact that the association between privative alleles and pollinators only occurs in epigenetic markers reflects the complexity of the relationship between genetic markers, epigenetic markers and propagule dispersion. Theoretical studies show that the adaptive capacity of a population can be induced or hindered by epigenetic mutation rate depending on the case and on whether these epimutations are environmentally biased or not (Greenspoon et al. 2022). In small populations however, adaptation to local conditions can be increased through epigenetic mutations even under a high migrant influx (Smithson et al. 2019). Our data add evidence on how epigenetic variability may behave differently to genetic variability particularly in small populations. Under the assumption that pollinator diversity increases pollen flow within and between populations, our findings suggests that due to the versatility of epigenetic mutations (Richards, 2006; Verhoeven et al. 2010; Johannes and Schmitz, 2019), and the fact that most species studied are long-lived, local epigenetic novelties in our narrow endemic species may remain unique in a scenario where there is insufficient pollen flow across populations to counteract population epigenetic differentiation. This highlights the contrasting sensitivity of epigenetic and genetic markers to pollen (and likely seed) flow and population dynamics, demonstrating the benefits of analysing both genetic and epigenetic markers in ecological studies focused in understanding how plant-pollinator interactions can modulate evolutionary processes in flowering plants and in restricted distribution species in particular.

## Conclusions

This study contributes to highlight the importance of pollination studies in understanding the evolution of plant rarity. First, the results corroborate findings from previous studies suggesting a lower pollinator visitation rate in plant species with a restricted distribution range in comparison to that in widespread congeners. Second, this study is the first in assessing the relationships between pollinator visitation rate and diversity and determinants of genetic and epigenetic diversity and distinctiveness in plant populations. Notably, the positive relationship of pollinators with genetic diversity and negative with epigenetic distinctiveness of populations of restricted species points to the importance of pollinators for plant species with small and isolated populations. In particular, gene flow and epigenetic novelties may be locally favourable and counterbalance genetic erosion in small populations inhabiting harsh environments.

## Acknowledgements

We thank all taxonomists who helped in the identification of insect specimens: Alejandro Núñez-Carbajal (Apoidea), Javier Ortiz (Halictidae), Tom Wood (Andrenidae) and Alberto Tinaut (Fomicidae). Teresa Boquete for assistance in field work. Pilar Bazaga and Esmeralda López-Perea for laboratory assistance. Consejería de Medio Ambiente, Junta de Andalucía, for authorizing this research. CA, CMH and MM conceived the ideas and designed methodology; CA, CMH, MM and JV collected the data; JV analysed the data and led the writing of the manuscript.

