## Appendix for "Assessing the links between pollinators and the genetic and epigenetic features of plant species with contrasting distribution ranges"

**Appendix S1.-** Pictures of the species considered in this study and spatial distribution of the sampled populations. Pictures on the left correspond to the widespread and those on the right to the restricted species. Blue symbols in the maps indicate the location of the corresponding widespread species and orange symbols those of the restricted congener.

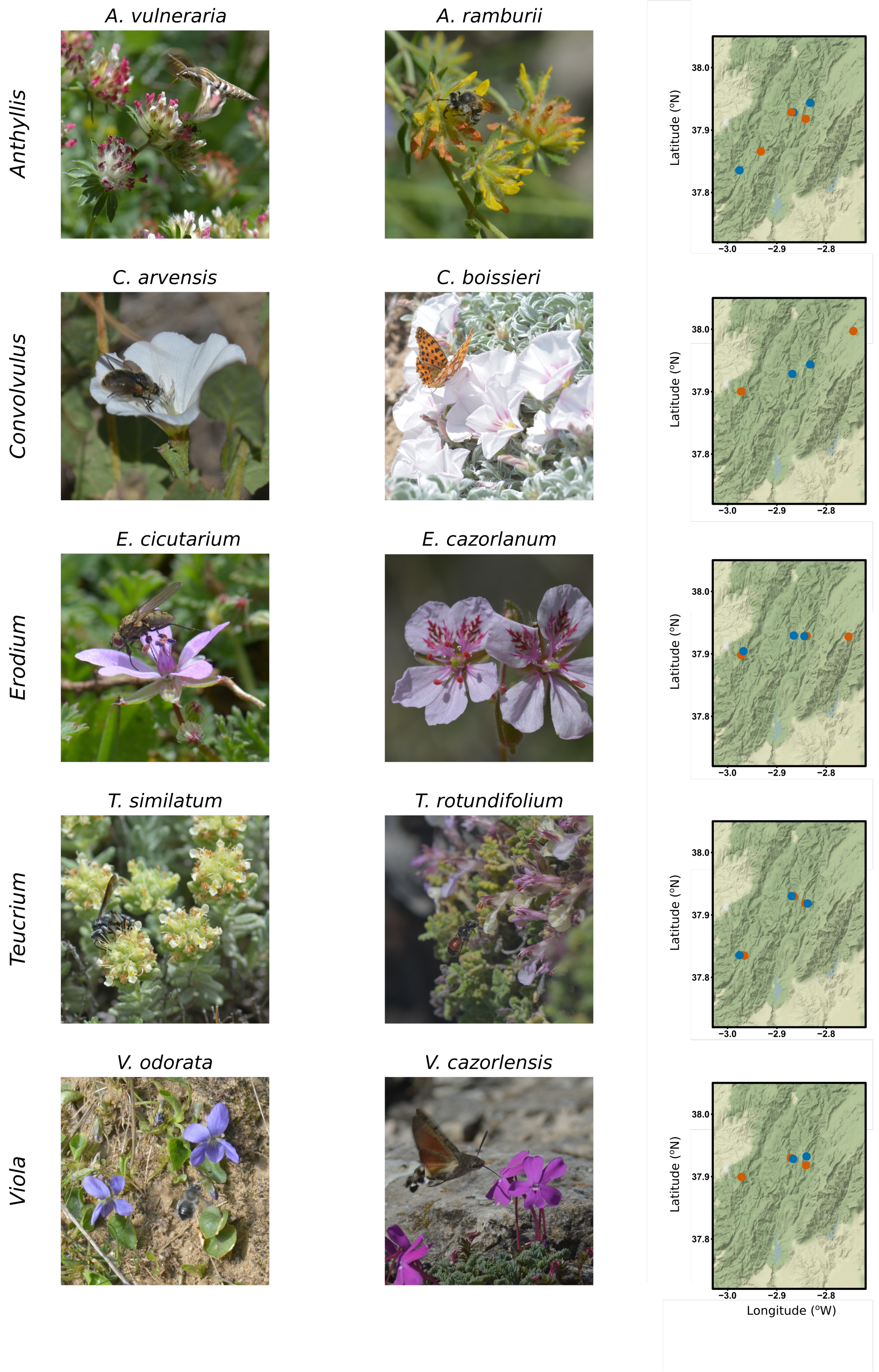

**Appendix S2.-** Sample species and populations. For each of the restricted (R) and widespread (W) studied species the description of the habitat type and distribution are specified. Details of the geographical location of each population are provided.

| Species | Habitat type | Distribution | Population | Latitude | Longitude | Altitude (masl) |
|---|---|---|---|---|---|---|
| *Anthyllis ramburii* (*R*) | High mountain dwarf procumbent shrub, dolomitic soils | Baetic ranges | anra1 | 37,92000 | -2,84150 | 1435 |
| | | | anra2 | 37,92842 | -2,86847 | 1624 |
| | | | anra3 | 37,86471 | -2,93298 | 1364 |
| *Anthyllis vulneraria* (*W*) | Dry grass, rocky environments with calcareous soils | Mediterranean basin | anvu1 | 37,84000 | -2,97576 | 1392 |
| | | | anvu3 | 37,94000 | -2,83258 | 1318 |
| *Convolvulus boissieri* (*R*) | High mountain dwarf procumbent shrub, dolomitic soils | Baetic ranges | cboi2 | 38,00000 | -2,74500 | 1728 |
| | | | cboi3 | 37,90039 | -2,97231 | 1521 |
| *Convolvulus arvensis* (*W*) | Cultivars, roadsides, wastelands | Temperate and tropical regions | carv1 | 37,93000 | -2,84129 | 1509 |
| | | | carv2 | 37,92982 | -2,86672 | 1632 |
| *Erodium cazorlanum* (*R*) | High mountain dry grass, rocky environments, dolomitic soils | Baetic ranges | ecazF | 37,93000 | -2,84140 | 1529 |
| | | | ecazL | 37,92772 | -2,75470 | 1755 |
| | | | ecazT | 37,89826 | -2,97258 | 1594 |
| *Erodium cicutarium* (*W*) | Meadows, flood plains, roadsides, gravel areas | Eurosiberian. Now circumpolar. | ecicC | 37,93000 | -2,86581 | 1628 |
| | | | ecicF | 37,90357 | -2,96959 | 1540 |
| | | | ecicT | 37,92886 | -2,84180 | 1518 |
| *Tecucrium rotundifolium* (*R*) | Rocky outcrops | Iberian Peninsula and Morocco | trot1 | 37,92000 | -2,84201 | 1468 |
| | | | trot2 | 37,83487 | -2,96609 | 1421 |
| | | | trot3 | 37,92972 | -2,86597 | 1639 |
| *Tecucrium similatum* (*W*) | Sclerophyllous and semideciduous forests. Calcareous scrublands and grasslands. Rocky mountain slopes | Iberian Peninsula | tpol1 | 37,84000 | -2,97587 | 1367 |
| | | | tpol2 | 37,91829 | -2,84106 | 1389 |
| | | | tpol3 | 37,93000 | -2,86582 | 1592 |
| *Viola cazorlensis* (*R*) | Rocky outcrops, cliffs, sandy dolomitic soils | Baetic ranges | vcaz1 | 37,92000 | -2,84074 | 1432 |
| | | | vcaz2 | 37,89960 | -2,97169 | 1518 |
| | | | vcaz3 | 37,93024 | -2,87163 | 1595 |
| *Viola odorata* (*W*) | Open woodlands, hedge banks and scrublands, edges of forests and clearings | Europe and Asia | vodo1 | 37,93000 | 2,84034 | 1458 |
| | | | vodo2 | 37,92878 | 2,86642 | 1629 |

**Appendix S3.-** Inter-annual stability of pollinator diversity

Herrera (2018) showed that contrary to the generalised decline in pollinator services in anthropised habitats, in well-conserved areas such as the Sierra de Cazorla this trend is more complex, being even neutral in many cases. To test the stability in pollinator diversity of our study populations we compared our data with that of Herrera (2018) and of other unpublished censuses.  We retrieved data from nine of our study populations from one or two different years: One population from A*nthyllis vulneraria*:  ANVU1 (years 2008 and 2015); One population from *Erodium cazorlanum*: ECAZF (2010), and one from *E. cicutarium*: ECICF (2015); One population from *Teucrium rotundifolium*: TROT3 (2007 and 2016), and one from *T. similatum*:  TPOL3 (2006, 2014); Two populations from *Viola cazorlensis*: VCAZ1 (year 2007), VCAZ2 (2015), and one from *V. odorata*: VODO1 (2010 and 2011). Data from our work and data from previous years where compared by using correlations on the observed species richness, Shannon and Simpson's diversities. These analyses showed a temporal stability in species richness while a more complex picture for the diversity measurements that take into consideration species abundances (see figure below).

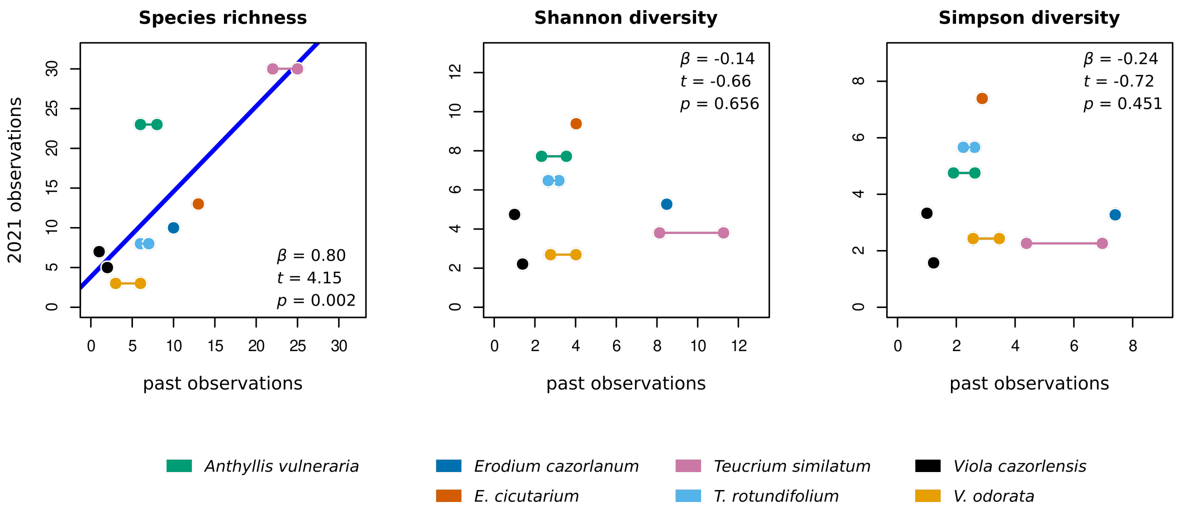


Figure caption: Connected dots depict data from the same population in different past years. The blue line in the left panel shows the linear regression of pollinator richness in the 2021 data against that of the data from past observations.

**Appendix S4.-** Checklist of pollinator working species. Each case corresponds to our working taxonomical unit. Thus, *'Species'* stands for the finest taxomical identification reached.

| Order | Family | 'Species' |
|---|---|---|
| Coleoptera | Anthicidae | Anthicidae sp1 |
| | Bruchidae | Bruchidae sp1 |
| | Bruchidae | Bruchidae sp2 |
| | Bruchidae | Bruchidae sp3 |
| | Bruchidae | Bruchidae sp4 |
| | Bruchidae | Bruchidae sp5 |
| | Buprestidae | *Anthaxia marmottani* |
| | Chrysomelidae | *Cryptocephalus ramburi* |
| | Coccinellidae | *Coccinella septempunctata* |
| | Coccinellidae | Coccinellidae sp. |
| | Coccinellidae | *Scymnus* sp. |
| | Curculionidae | *Cycloderes* sp. |
| | Curculionidae | *Miarus campanulae* |
| | Dermestidae | *Anthrenus angustefasciatus* |
| | Dermestidae | *Anthrenus festivus* |
| | Dermestidae | *Attagenus trifasciatus* |
| | Dermestidae | *Orphilus niger* |
| | Meloidae | *Mylabris* sp. |
| | Melyridae | *Dasytes terminalis* |
| | Mordellidae | *Mordella* sp. |
| | Nitidulidae | *Brassicogethes aeneus* |
| | Nitidulidae | Nitidulidae |
| | Oedemeridae | *Chrysanthia viridissima* |
| | Oedemeridae | *Oedemera* sp. |
| | Tenebrionidae | *Heliotaurus ruficolis* |
| Diptera | Anthomyiidae | Anthomyiidae sp1 |
| | Anthomyiidae | Anthomyiidae sp2 |
| | Anthomyiidae | Anthomyiidae sp3 |
| | Anthomyiidae | Anthomyiidae sp4 |
| | Anthomyiidae | Anthomyiidae sp5 |
| | Anthomyiidae | Anthomyiidae spp. |
| | Bibionidae | Bibionidae |
| | Bibionidae | *Dilophus* sp. |
| | Bombyliidae | *Bombylius cruciatus* |
| | Bombyliidae | *Bombylius fimbriatus* |
| | Bombyliidae | *Bombylius major* |
| | Bombyliidae | *Bombylius* sp. |
| | Bombyliidae | *Bombylius venosus* |
| | Bombyliidae | *Hemipenthes morio* |
| | Bombyliidae | *Phthiria* sp. |
| | Bombyliidae | *Systoechus* sp1 |
| | Bombyliidae | *Systoechus* sp2 |
| | Bombyliidae | *Usia pubera* |
| | Bombyliidae | *Usia* sp. |
| | Bombyliidae | *Villa* sp. |
| | Calliphoridae | *Rhyncomyia felina* |
| | Chloropidae | *Tricimba cincta* |
| | Conopidae | *Myopa* sp. |
| Diptera | Empididae | *Empis* sp. |
| | Empididae | *Hilara* sp. |
| | Empididae | *Ramphomyia* sp. |
| | Heleomyzidae | *Suillia* sp. |
| | Muscidae | *Neomyia* sp. |
| | Mythicomyiidae | Mythicomyiidae |
| | Sarcophagidae | Sarcophagidae |
| | Syrphidae | *Chrysotoxum intermedium* |
| | Syrphidae | *Eristalis tenax* |
| | Syrphidae | *Eupeodes corollae* |
| | Syrphidae | *Melanostoma scalare* |
| | Syrphidae | *Paragus albifrons* |
| | Syrphidae | *Pelecocera* sp. |
| | Syrphidae | *Scaeva pyrastri* |
| | Syrphidae | *Sphaerophoria scripta* |
| | Tachinidae | *Peleteria varia* |
| | Tachinidae | *Prosena siberita* |
| | Tachinidae | *Rhamphina rectirostris* |
| | Tachinidae | Tachinidae sp1 |
| | Tephritidae | *Campiglossa* sp. |
| | Tephritidae | Tephritidae |
| Hemiptera | Miridae | *Hadrodemus m-flavum* |
| | Pentatomidae | *Dolycoris baccarum* |
| | Rhopalidae | Rhopalidae |
| | | Hemiptera sp1 |
| | | Hemiptera sp2 |
| | | Hemiptera sp3 |
| Hymenoptera | Andrenidae | *Andrena asperrima* |
| | Andrenidae | *Andrena* sp1 |
| | Andrenidae | *Andrena* sp2 |
| | Andrenidae | *Andrena* sp3 |
| | Andrenidae | *Andrena* sp5 |
| | Andrenidae | *Andrena* sp6 |
| | Andrenidae | *Andrena ferrugineicrus* |
| | Andrenidae | *Andrena granulosa* |
| | Andrenidae | *Andrena icterina* |
| | Andrenidae | *Andrena labialis* |
| | Andrenidae | *Andrena labiata* |
| | Andrenidae | *Andrena nigroaenea* |
| | Andrenidae | *Andrena ovatula* |
| | Andrenidae | *Andrena pilipes* |
| | Andrenidae | *Andrena schenki* |
| | Andrenidae | *Andrena similis* |
| | Andrenidae | *Andrena* sp. |
| | Apidae | *Anthophora aestivalis* |
| | Apidae | *Anthophora atroalba* |
| | Apidae | *Anthophora dispar* |
| | Apidae | *Anthophora plumipes* |

**Appendix S4 (cont.).-** Checklist of pollinator working species.

| Order | Family | 'Species' |
|---|---|---|
| Hymenoptera | Apidae | *Anthophora retusa* |
| | Apidae | *Anthophora* sp. |
| | Apidae | *Apis mellifera* |
| | Apidae | *Bombus pratorum* |
| | Apidae | *Bombus* sp. |
| | Apidae | *Bombus terrestris* |
| | Apidae | *Ceratina chalybea* |
| | Apidae | *Ceratina* sp. |
| | Apidae | *Eucera caspica* |
| | Apidae | *Eucera* sp1 |
| | Apidae | *Eucera* sp2 |
| | Apidae | *Nomada* sp. |
| | Apidae | *Xylocopa cantabrita* |
| | Braconidae | Agathidinae |
| | Braconidae | Braconinae |
| | Chrysididae | Chrysididae |
| | Crabronidae | *Cerceris* sp. |
| | Formicidae | *Formica cuniculata* |
| | Formicidae | Formicidae sp. |
| | Formicidae | *Iberoformica subrufa* |
| | Formicidae | *Plagiolepis pygmaea* |
| | Formicidae | *Proformica longiseta* |
| | Formicidae | *Tapinoma madeirense* |
| | Halictidae | Halictidae |
| | Halictidae | *Halictus crenicornis* |
| | Halictidae | *Halictus* sp1 |
| | Halictidae | *Halictus subauratus* |
| | Halictidae | *Halictus tridivisus* |
| | Halictidae | *Lasioglossum aeratum* |
| | Halictidae | *Lasioglossum buccale* |
| | Halictidae | *Lasioglossum* sp1 |
| | Halictidae | *Lasioglossum* sp2 |
| | Halictidae | *Lasioglossum* sp5 |
| | Halictidae | *Lasioglossum laevigatum* |
| | Halictidae | *Lasioglossum malachurum* |
| | Halictidae | *Lasioglossum marginatum* |
| | Halictidae | *Lasioglossum minutissimum* |
| | Halictidae | *Lasioglossum nitidiusculum* |
| Hymenoptera | Halictidae | *Lasioglossum nitidulum* |
| | Halictidae | *Lasioglossum pauperatum* |
| | Halictidae | *Lasioglossum* sp. |
| | Halictidae | *Lasioglossum subaenescens* |
| | Halictidae | *Lasioglossum subhirtum* |
| | Halictidae | *Seladonia* sp. |
| | Megachilidae | *Megachile pilidens* |
| | Megachilidae | Megachilidae |
| | Megachilidae | *Osmia andrenoides* |
| | Megachilidae | *Osmia* sp1 |
| | Megachilidae | *Osmia* sp2 |
| | Megachilidae | *Osmia submicans* |
| | Megachilidae | *Osmia tricornis* |
| | Megachilidae | *Protosmia* sp. |
| | Melittidae | *Melitta* sp. |
| Lepidoptera | Choreutidae | *Prochoreutis* sp. |
| | Hesperiidae | *Carcharodus baeticus* |
| | Hesperiidae | *Pyrgus malvoides* |
| | Hesperiidae | *Thymelicus sylvestris* |
| | Lycaenidae | *Lycaena phlaeas* |
| | Lycaenidae | *Polyommatus bellargus* |
| | Lycaenidae | *Polyommatus icarus* |
| | Lycaenidae | *Satyrium spini* |
| | Nymphalidae | *Argynnis niobe* |
| | Nymphalidae | *Issoria lathonia* |
| | Nymphalidae | *Lasiommata megera* |
| | Nymphalidae | *Vanessa cardui* |
| | Pieridae | *Colias croceus* |
| | Pieridae | *Pieris napi* |
| | Pterophoridae | Pterophoridae |
| | Scythrididae | *Scythris* sp. |
| | Sphingidae | *Macroglossum stellatarum* |
| Orthoptera | Acrididae | Acrididae |
| | Acrididae | *Calliptamus* sp. |
| | Acrididae | *Pezotettix giornae* |
| | Tettigoniidae | Tettigoniinae sp1 |
| | Tettigoniidae | Tettigoniinae sp2 |

**Appendix S5.-** Genetic and epigenetic data

Plant samples were obtained from newly expanded, fully-grown leaves to avoid intra-individual variation in methylation levels.

The AFLP analysis was performed using standard protocols involving the use of fluorescent dye-labeled selective primers. For each genus four combinations of the same MseI+3 / PstI+2 selective primer pairs were used providing reliable and consistently scorable results. All primer combinations used for each species were merged in a raw single binary genetic data matrix indicating the presence or absence of each AFLP fragment.

The MSAP technique combines a frequent cutter restriction enzyme (here we used MseI) with two methylation-sensitive restriction enzymes (HpaII and MspI) in parallel runs. HpaII and MspI recognize and cleave the same sequence, 5′-CCGG, but differ in their sensitivity to the methylation state of the cytosines. Both enzymes cut the DNA if the restriction site is not methylated, but whereas MspI cuts sites with one or two methylated internal cytosines, HpaII only cuts sites with one methylated external cytosine. For each genus we used three to four MseI+3 / HpaII–MspI+2 selective primer pair combinations. We followed Schultz *et al*., (2014) to analyse the MSAP profiles, obtaining two raw binary epigenetic data matrices: one with hemi- or fully methylated epiloci (hereafter M-MSAP), and one data matrix of unmethylated epiloci (U-MSAP). We eliminated inconsistent loci after measuring their repeatability in a subset of samples (8.6–17.6 % of samples for AFLP; 13.3–29.3% of samples for MSAP). Moreover, we also discarded non-informative monomorphic loci by selecting those in which the proportion of presence or absence of the fragment exceeded 5%. This yielded 140 to 316 AFLP, 84 to 181 U-MSAP and 100 to 213 M-MSAP loci per species.

**Appendix S6.-** Description of some of the diversity and distinctiveness indices used.

**a) Per-locus average of the Shannon's diversity index** (SI):

This index is derived from the Shannon-Weaver index (Shannon, 1948) and is defined as:

$$SI=\frac{-\sum_{i=1}^{n} P_i \cdot \log_e(P_i)}{n}$$

where the formula within the summation in the numerator corresponds to the Shannon's Index of Phenotypic Diversity (King and Schaal, 1989) for a given marker. The number of markers is given by *n* and the frequency of the presence of the *i*th marker by $P_i$.

**b) Rarity Index** (*RI*):

This index is an adaptation to population genetics of the range-down-weighted species values used in biogeographical research (e.g. Crisp *et al*., 2001). It is defined as:

$$RI_x=\sum_{i=1}^{n} \frac{S_{ix}}{\sum_{j=1}^{k} S_{ij}}$$

where *n* is the number of markers, $S_{ij}$ denotes the binary state of the *i*th marker (present 1 or absent 0) in a given individual *x*. The total number of individuals in the population is indicated by *k*. Due to the accumulation of rare alleles in isolated populations over a long period of time, the index value is expected to be high in populations with an old vicariance.

**Appendix S7.-** Model selection for the relationships of pollinators with population genetic and epigenetic diversity and distinctiveness of plant species with restricted distribution range. Each model selection is based on previous significant correlations. Models are ranked based on the Akaike Information Criterion (AIC) and on the comparisons between models, such that the chosen model is on top. Parameters reported were calculated using the Maximum Likelihood criterion. *P*-values in bold denote significance at α = 0.05. Note that for the proportion of privative methylated epialleles (M-MSAP) models use a log-link function. *R* syntaxes of each model are depicted in the table's foot notes.

*Genetic and epigenetic diversity*

| Response | Predictor | Random effects structure | AIC | BIC | Bayes factor | *P* |
|---|---|---|---|---|---|---|
| AFLP *Shannon diversity* | Patch visitation probability | Intercept | -46.47 | -43.91 | 0.86 | 0.036 |
| | | Intercept and slope * | -48.04 | -44.21 | 1.16 | |
| | Flower visitation probability | Intercept | -44.06 | -41.50 | 1.64 | 0.066 |
| | | Intercept and slope | -44.35 | -40.52 | 0.61 | |
| | $^{0}D$ | Intercept | -36.38 | -33.83 | 6.31 | 0.533 |
| | | Intercept and slope * | -33.98 | -30.14 | 0.16 | |
| U-MSAP *Shannon diversity* | Patch visitation probability | Intercept | -40.36 | -37.80 | 11.89 | 0.813 |
| | | Intercept and slope * | -36.68 | -32.85 | 0.08 | |
| | $^{0}D$ | Intercept | -34.35 | -31.79 | 10.70 | 0.887 |
| | | Intercept and slope | -30.89 | -27.05 | 0.09 | |
| | $^{1}D$ | Intercept | -34.53 | -31.97 | 8.96 | 0.832 |
| | | Intercept and slope | -31.42 | -27.59 | 0.11 | |

*Genetic and epigenetic distinctiveness*

| Response | Predictor | Random effects structure | AIC | BIC | Bayes factor | *P* |
|---|---|---|---|---|---|---|
| AFLP *Rarity index* | Patch visitation probability | Intercept | 29.06 | 31.62 | 2.10 | 0.230 |
| | | Intercept and slope | 29.26 | 33.10 | 0.48 | |
| M-MSAP *Prop. of privative alleles* (log-link function) | $^{0}D$ | Intercept | -58.76 | -56.21 | 0.03 | 0.002 |
| | | Intercept and slope † | -67.08 | -63.25 | 33.86 | |
| | $^{1}D$ | Intercept | -58.85 | -56.29 | 0.08 | 0.005 |
| | | Intercept and slope † | -65.32 | -61.49 | 13.41 | |
| | $^{2}D$ | Intercept | -58.17 | -55.61 | 0.60 | 0.043 |
| | | Intercept and slope * | -60.48 | ~~56~~64 | 1.67 | |

***** singular fit; **†** convergence failure
Random intercept models: *lmer(Y ~ X + (1 | genus))*
Random intercept and slope models: *lmer(Y ~ X + (X | genus))*
Where *Y* and *X* denote the response and predictor variables respectively. M-MSAP models use generalized mixed effects models (*glmer*) and log-link function (*family = gaussian(link="log")*).

**Appendix S8.-** Rank-abundance distributions of pollinator species at each population. *W* denotes widespread species, *R* indicates restricted species. Colour codes denote the Order of each pollinator species. Plots are order in genus. Panels are sorted by plant genus.

### a) *Anthyllis*

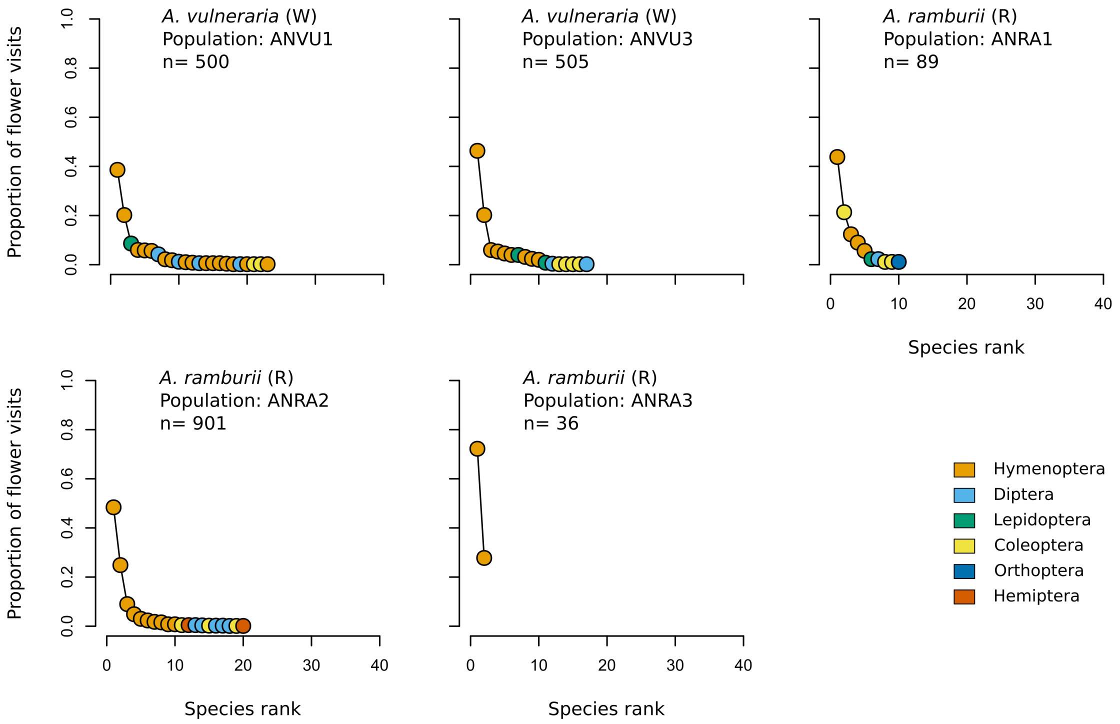


### b) *Convolvulus*

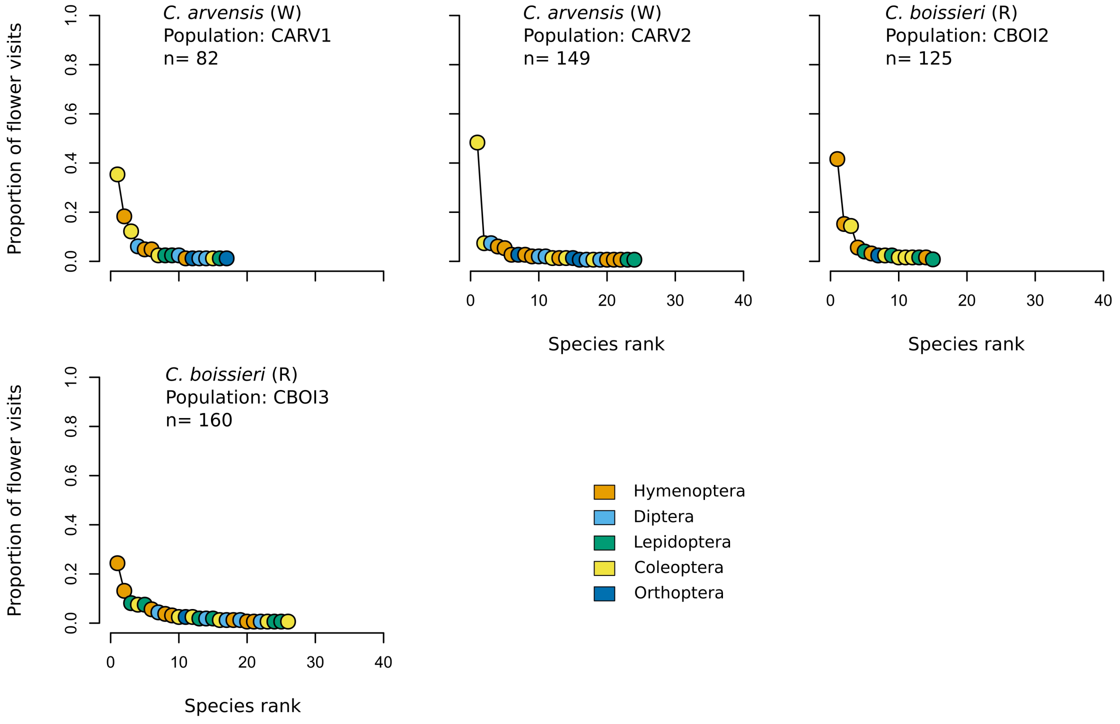

### c) *Erodium*

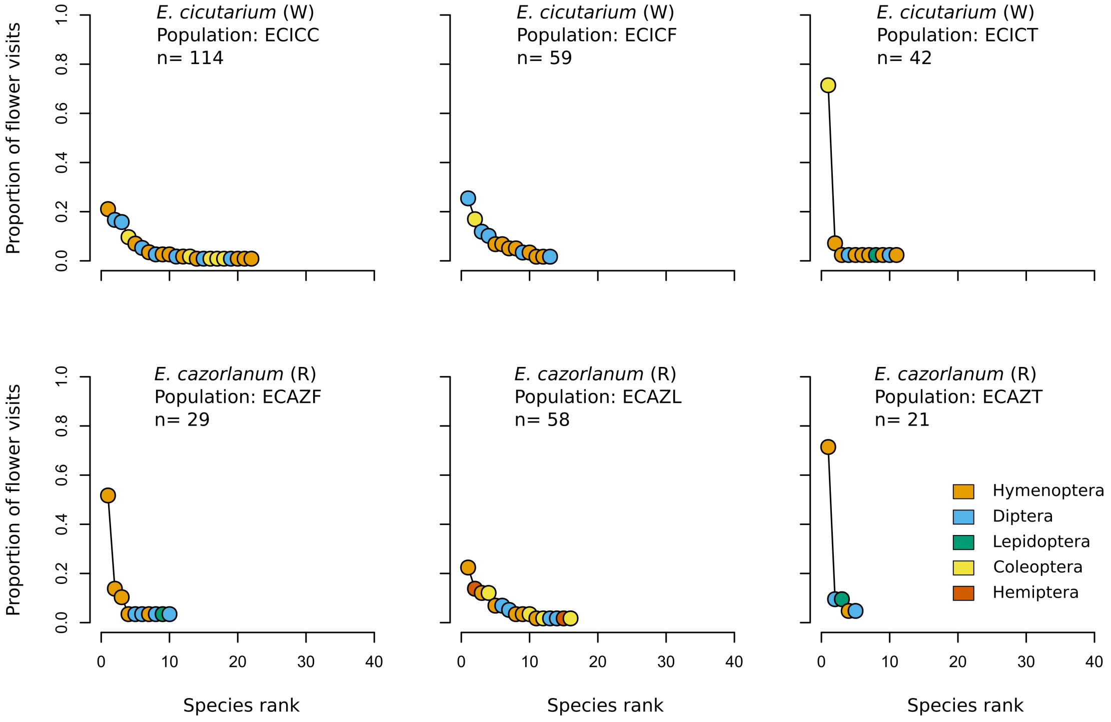


### d) *Teucrium*

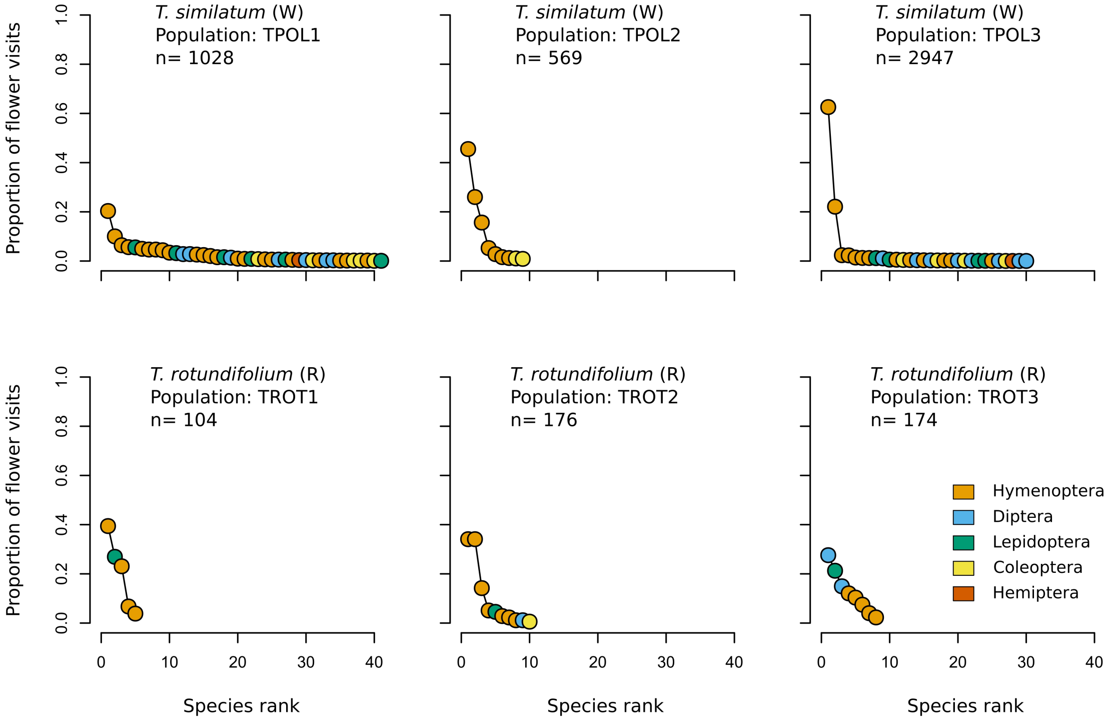

### c) ***Viola***

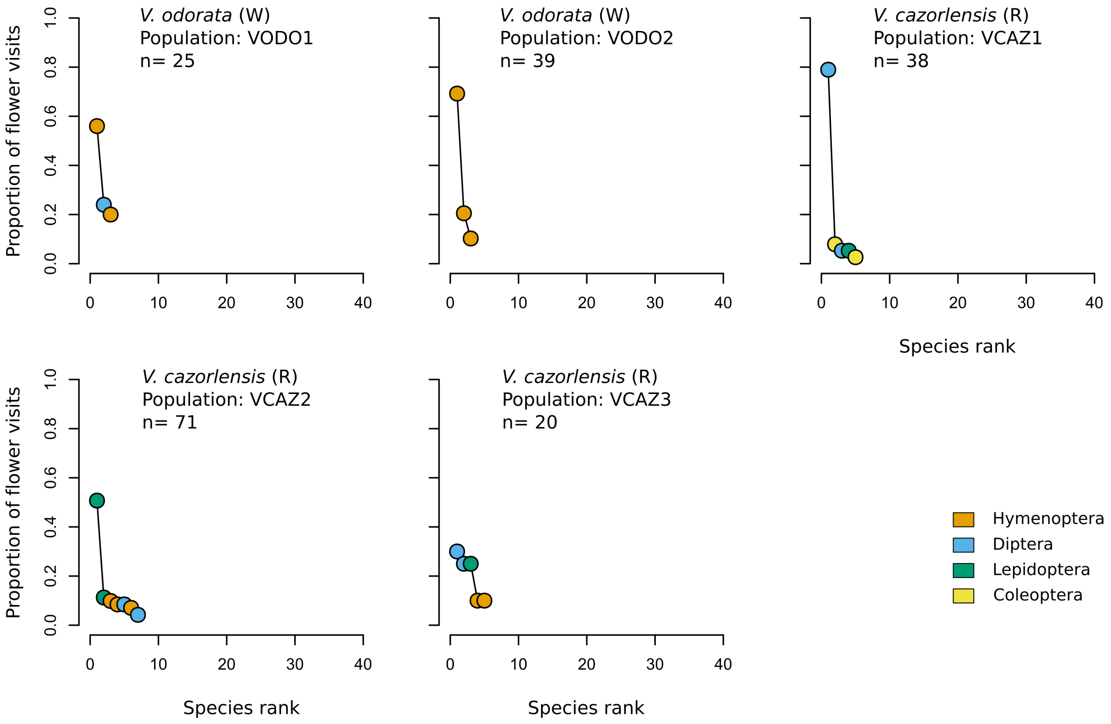
V. odorata (W)
Population: VODO1
n= 25
V. odorata (W)
Population: VODO2
n= 39
V. cazorlensis (R)
Population: VCAZ1
n= 38
V. cazorlensis (R)
Population: VCAZ2
n= 71
V. cazorlensis (R)
Population: VCAZ3
n= 20
Proportion of flower visits
Species rank
Hymenoptera
Diptera
Lepidoptera
Coleoptera

**Appendix S9.-** Pollinator diversity profiles. Panels show the diversity profiles of the sampled populations from congeneric widespread (open symbols) and restricted species (filled symbols) at the flower level. Diversity values correspond to the Hill numbers (${}^{q}D$) calculated for different values of $q$ at a sampling coverage of 0.8. Bigger symbols highlight Diversity values corresponding to species richness (q=0); logarithm of the Shannon diversity (q=1) and Simpson diversity (q=2).

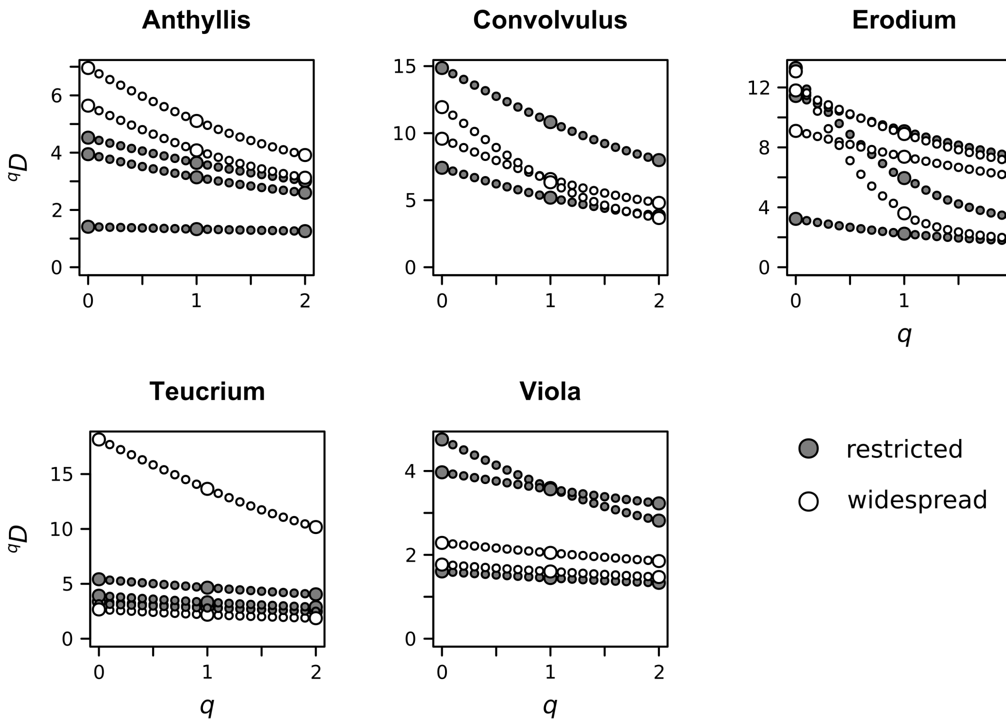

**Appendix S10.-** Results of the linear mixed models examining the effect of the type of distribution (restricted or widespread) on the patch and flower visitation probabilities. For each predictor, its estimate, *z* statistic and *P* value are shown. Note that at the patch level the number of flowers per patch is included as an additional predictor. Parameters of the random effects include the residual variance across samples ($\sigma^2$) and the variance attributed to each of the levels ($\tau$). Significant *P* values at $\alpha = 0.05$ are highlighted in bold. *R* syntax of models is depicted in the table's foot notes.

| | Patch visitation probability | | | Flower visitation probability | | |
|---|---|---|---|---|---|---|
| | Estimate | *z* | *P* | Estimate | *z* | *P* |
| Intercept | -0.13 | -0.30 | 0.761 | -4.18 | -7.45 | **<0.001** |
| Distribution (restricted) | -0.73 | -2.09 | **0.037** | -0.73 | -1.95 | 0.051 |
| No. flowers | 0.33 | 5.37 | **<0.001** | | | |
| Random effects: | | | | | | |
| $\sigma^2$ | 3.29 | | | 8.11 | | |
| $\tau$ | 0.58 (genus) ; 0.69 (population) | | | 1.19 (genus) ; 0.82 (population) ; 4.82 (census) | | |
| Variance explained: | | | | | | |
| | $R^2_{MARGINAL} = 0.050$<br>$R^2_{CONDITIONAL} = 0.314$ | | | $R^2_{MARGINAL} = 0.013$<br>$R^2_{CONDITIONAL} = 0.209$ | | |

**Patch visitation probability model:**
*glmer(Y ~ distr + display + (1 | genus / pop), family = binomial(link = "logit"))*
where *Y* is a binomial response of whether a patch has been visited (*Y*=1) or not (*Y*=0), *distr* is the distribution type (restricted or widespread), *display* is a scaled covariate of the number of flowers at each patch, and *pop* denotes the population.
**Flower visitation probability model:**
*glmer(cbind(Y1, Y0) ~ distr + (1 | genus / pop / ID), family = binomial(link = "logit"))*
where *Y1* and *Y0* respectively denote the number of contacted and not contacted flowers in the patch, and *ID* identifies each census.

**Appendix S11.-** Post-hoc paired comparisons of patch and flower visitation probabilities between congeneric pairs of widespread and restricted species. For each comparison the estimated odds ratio and its standard error, in parenthesis, are are shown. Significant *P* values at $\alpha = 0.05$ are highlighted in bold.

| Pair comparison (widespread / restricted) | Patch visitation probability | | | Flower visitation probability | | |
|---|---|---|---|---|---|---|
| | Odds ratio | *z* ratio | *P* value | Odds ratio | *z* ratio | *P* value |
| *Anthyllis vulneraria* / *A. ramburii* | 2.547 (1.405) | 1.694 | 0.090 | 2.190 (0.874) | 1.965 | **0.049** |
| *Convolvulus arvensis* / *C. boissieri* | 1.124 (0.676) | 0.194 | 0.847 | 5.130 (2.249) | 3.729 | **<0.001** |
| *Erodium cicutarium* / *cazorlanum* | 1.490 (0.736) | 0.808 | 0.419 | 0.871 (0.381) | -0.316 | 0.752 |
| *Teucrium similatum* / *rotundifolium* | 9.048 (4.659) | 4.277 | **<0.001** | 10.774 (4.212) | 6.081 | **<0.001** |
| *Viola odorata* / *V. cazorlensis* | 0.529 (0.315) | -1.070 | 0.285 | 0.556 (0.251) | -1.300 | 0.194 |

**Appendix S12.-** Correlations between pollinator and genetic and epigenetic diversity and distinctiveness. The panels show the correlation values between each pair of indices calculated for AFLP, U-MSAP and M-MSAP markers. Values for the restricted species are characterised by circles, while squares depict values for the widespread species. Filled symbols indicate significant correlations at α = 0.05. Increased size symbols highlight correlations at $^{0}D$ (species richness), $^{1}D$ (logarithm of the Shannon diversity index) and $^{2}D$ (Simpson diversity index).

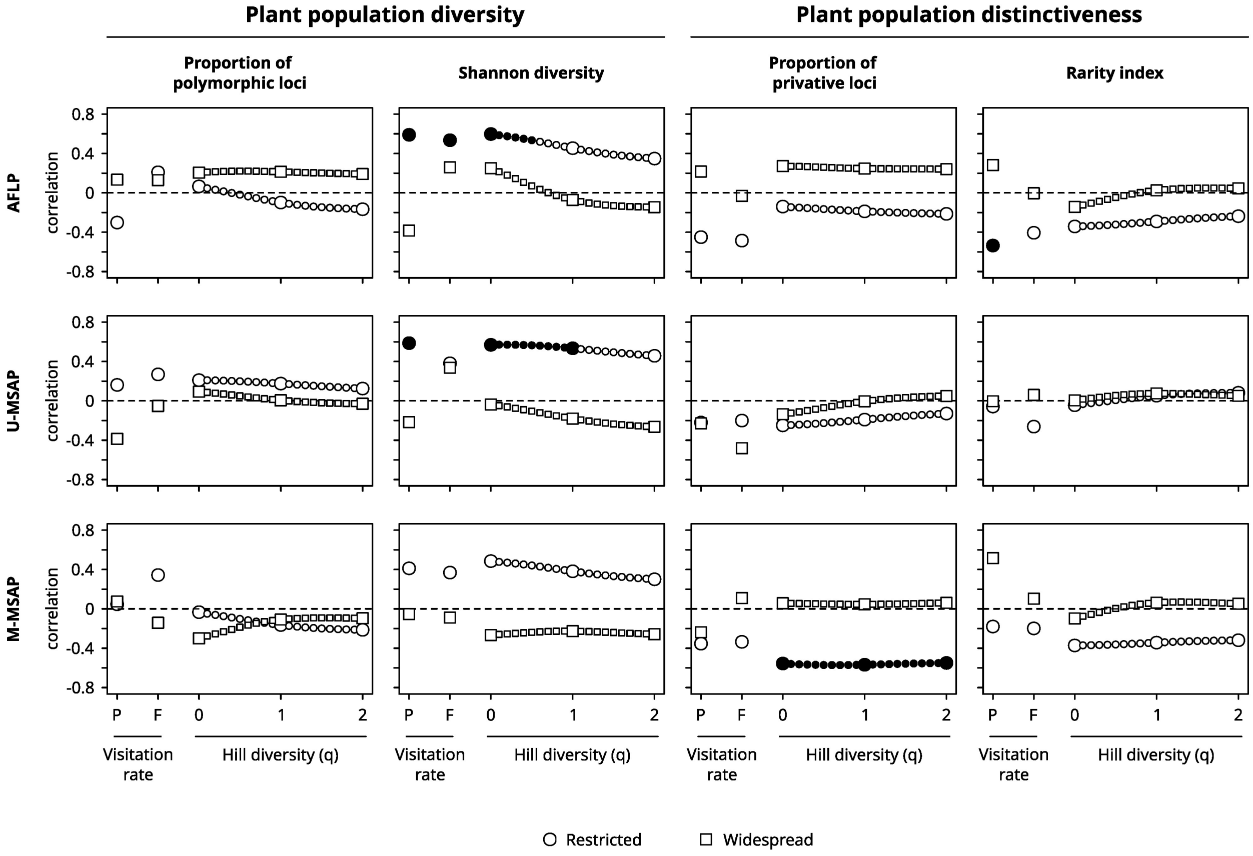
